# Domain-specific quantification of prion protein in cerebrospinal fluid by targeted mass spectrometry

**DOI:** 10.1101/591487

**Authors:** Eric Vallabh Minikel, Eric Kuhn, Alexandra R Cocco, Sonia M Vallabh, Christina R Hartigan, Andrew G Reidenbach, Jiri G Safar, Gregory J Raymond, Michael D McCarthy, Rhonda O’Keefe, Franc Llorens, Inga Zerr, Sabina Capellari, Piero Parchi, Stuart L Schreiber, Steven A Carr

## Abstract

Therapies currently in preclinical development for prion disease seek to lower prion protein (PrP) expression in the brain. Trials of such therapies are likely to rely on quantification of PrP in cerebrospinal fluid (CSF) as a pharmacodynamic biomarker and possibly as a trial endpoint. Studies using PrP ELISA kits have reproducibly shown that CSF PrP is lowered in the symptomatic phase of disease, a potential confounder for reading out the effect of PrP-lowering drugs in symptomatic patients. To date it has been unclear whether the reduced abundance of PrP in CSF results from its incorporation into plaques, retention in intracellular compartments, downregulation as a function of the disease process, or other factors. Because misfolding or proteolytic cleavage could potentially render PrP invisible to ELISA even if its concentration were constant or increasing in disease, we sought to establish an orthogonal method for CSF PrP quantification. We developed a targeted mass spectrometry method based on multiple reaction monitoring (MRM) of nine PrP tryptic peptides quantified relative to known concentrations of isotopically labeled standards. Analytical validation experiments showed process replicate coefficients of variation below 15%, good dilution linearity and recovery, and suitable performance for both CSF and brain homogenate and across humans as well as preclinical species of interest. In *N*=55 CSF samples from individuals referred to prion surveillance centers with rapidly progressive dementia, all six human PrP peptides, spanning the N- and C-terminal domains of PrP, were uniformly reduced in prion disease cases compared to individuals with non-prion diagnoses. This confirms the findings from ELISA studies, demonstrating that lowered CSF PrP concentration in prion disease is a genuine result of the disease process and not merely an artifact of ELISA-based measurement. We provide a targeted mass spectrometry-based method suitable for preclinical and clinical quantification of CSF PrP as a tool for drug development.

## Introduction

Prion disease is a fatal and incurable neurodegenerative disease caused by misfolding of the prion protein (PrP), and may be sporadic, genetic, or acquired^1^. Therapies currently in preclinical development for prion disease seek to lower PrP levels in the brain, a genetically well-validated strategy^2^. Clinical trials of PrP-lowering agents will rely on quantification of PrP in cerebrospinal fluid (CSF) as, at a minimum, a pharmacodynamic biomarker^3^. This marker may, however, have even greater importance. Predictive testing of pre-symptomatic individuals harboring highly penetrant genetic mutations^4^ that cause prion disease provides an opportunity for early therapeutic intervention to preserve healthy life, but randomization to a clinical endpoint in this population appears infeasible^5^. The U.S. Food and Drug Administration has indicated its willingness to consider lowered CSF PrP in this population as a potential surrogate endpoint for Accelerated Approval^2,6^. Precise quantification of PrP in CSF will be essential to the development of prion disease therapeutics.

PrP is an extracellular GPI-anchored protein that can be shed from the plasma membrane by ADAM10 and other peptidases^7,8^. CSF PrP is predominantly soluble and full-length^9^, suggesting that it originates chiefly from this proteolytic shedding near the C terminus, although lower molecular weight fragments of PrP have also been identified in CSF^10^, which may originate from other endoproteolytic events^7,11^, and anchored PrP is also released from cells on exosomes^12^. PrP is sufficiently abundant in CSF, at concentrations of tens or hundreds of nanograms per milliliter, to be readily quantified with enzyme-linked immunosorbent assay (ELISA). Studies using ELISA have reproducibly found that CSF PrP is decreased in the symptomatic phase of prion disease^3,13–16^. Therefore, even though CSF PrP is brain-derived and exhibits good within-subject test-retest reliability in individuals without prion disease^3^, it might be difficult to use this biomarker to read out the effect of a PrP-lowering drug in symptomatic individuals, because it is unclear whether to expect that such a drug should cause a further decrease in CSF PrP as a direct pharmacodynamic effect, or an increase in CSF PrP due to alleviation of the disease process. This confounder could potentially limit the use of ELISA-based CSF PrP quantification as a pharmacodynamic biomarker to pre-symptomatic individuals only.

Prion disease is caused by a gain of function^1^, and animal studies have shown that total PrP in the brain increases over the course of prion disease as misfolded PrP accumulates^17–19^. The paradoxical decrease in PrP in CSF during prion disease might be due to its incorporation into plaques^20^, diversion into intracellular locations^21,22^, or downregulation as a function of the disease process^23^. However, occlusion of epitopes due to misfolding^24^ or upregulation of proteolytic cleavage in disease^7,23,25^ could also render PrP invisible to ELISA even if its concentration were constant or increasing. We therefore sought to establish an orthogonal method for CSF PrP quantification.

Here, we describe quantification of CSF PrP using a form of targeted mass spectrometry — multiple reaction monitoring (MRM)^26^. We analyze *N*=55 clinical samples from prion and non-prion disease patients by PrP MRM and find that six out of six PrP tryptic peptides, spanning N- and C-terminal domains of the protein, are uniformly decreased in prion disease. Thus, PrP concentration is genuinely lowered in prion disease CSF. Our findings supply an alternative method for validating the findings of ELISA-based studies of CSF PrP, and provide a potential assay for use as a pharmacodynamic biomarker in preclinical drug development and in human trials.

## Methods

### Cerebrospinal fluid and brain samples

This study was approved by the Broad Institute’s Office of Research Subjects Protection (ORSP-3587). Written consent for research use of samples was obtained from patients or next of kin as appropriate.

All CSF samples in this study have been previously reported^3^. CSF samples for assay development were large volume normal pressure hydrocephalus samples provided by MIND Tissue Bank at Massachusetts General Hospital. Clinical CSF samples were premortem lumbar punctures from individuals referred to prion surveillance centers in Italy (Bologna) or Germany (Göttingen) with suspected prion disease and who were later either determined by autopsy or probable diagnostic criteria^27^ including real-time quaking-induced conversion (RT-QuIC^28^) as prion disease, or confirmed as non-prion cases on the basis of autopsy, patient recovery, or definitive other diagnostic test. Individuals with non-prion diagnoses (*N*=21) included autoimmune disease (*N*=8), non-prion neurodegenerative disease (*N*=6), psychiatric illness (*N*=3), stroke (*N*=1), brain cancer (*N*=1), and other (*N*=2). Sporadic prion disease cases (*N*=23) included probable cases (*N*=10) and autopsy-confirmed definite cases (*N*=13, of subtypes: 6 MM1, 3 VV2 and 4 other/unknown). Genetic prion disease cases (*N*=11) included D178N (*N*=2), E200K (*N*=7), and V210I (*N*=2). Samples were de-identified and broken into five batches (to be run on different days) randomly using an R script. Assay operators were blinded to diagnosis. PrP ELISA, hemoglobin, and total protein measurements on these CSF samples were previously reported^3^.

Rat and cynomolgus monkey CSF were purchased from BioIVT. Human brain tissue was from a non-prion disease control individual provided by the National Prion Disease Pathology Surveillance Center (Cleveland, OH). Mouse brain tissue from Edinburgh PrP knockout mice^29^ backcrossed to a C57BL/10 background^30^, and matching tissue from wild-type C57BL/10 mice, were provided by Gregory J. Raymond (NIAID Rocky Mountain Labs, Hamilton, MT).

### Recombinant protein preparation

Untagged recombinant HuPrP23-230 (MW=22,878) and MoPrP23-231 (MW=23,151), corresponding to full-length post-translationally modified human and mouse PrP without the signal peptide or GPI signal but retaining an N-terminal methionine, were purified by denaturation and Ni-NTA affinity from *E. coli* inclusion bodies as previously described^31,32^, using a vector generously provided by Byron Caughey (NIAID Rocky Mountain Labs, Hamilton, MT). ^15^N incorporation was achieved by growing the *E. coli* in ^15^N cell growth medium (Cambridge Isotope Laboratories CGM-1000-N) induced with ^15^N auto-induction medium (Millipore 71759-3). Protein concentration was determined by amino acid analysis (AAA, New England Peptide). Percent ^15^N isotopic incorporation was estimated using LC-MS/MS. ^15^N labeled human recombinant prion protein (10 µg) was digested and desalted following the procedure as described in *PrP MRM assay* and analyzed as described in *Pilot LC-MS/MS analysis*. Precursor masses for ^15^N were extracted from the chromatograms using XCalibur software Qualbrowser software (Thermo) 3.0.63 with a 6 mz window of centered on the precursors and charge states listed in Table S1. Isotopic envelopes between protein expressed in 15N containing media and standard media were compared visually. Summation of all observed mz peak areas less than the ^12^C monoisotopic mass peak were compared to summation of all expected isotope peak to estimate the overall completeness of ^15^N incorporation (Figure S1).

### Pilot LC-MS/MS analyses of CSF and recombinant PrP

Samples of dried digested recombinant proteins or human cerebrospinal fluid (processed as described in *PrP MRM assay*) were reconstituted in 3% acetonitrile/5% acetic acid to a final concentration of approximately 1 µg total protein per 1 µL and analyzed in a single injection using a standard 2h reversed-phase gradient. LC-MS/MS was performed using a QExactive mass spectrometer (Thermo) equipped with a Proxeon Easy-nLC 1200 and a custom built nanospray source (James A. Hill Instrument Services). Samples were injected (1 to 2 µg) onto a 75 um ID PicoFrit column (New Objective) packed to 20 cm with Reprosil-Pur C18 AQ 1.9 um media (Dr. Maisch) and heated to 50°C. MS source conditions were set as follows: spray voltage 2000, capillary temperature 250, S-lens RF level 50. A single Orbitrap MS scan from 300 to 1800 m/z at a resolution of 70,000 with AGC set at 3e6 was followed by up to 12 MS/MS scans at a resolution of 17,500 with AGC set at 5e4. MS/MS spectra were collected with normalized collision energy of 25 and isolation width of 2.5 amu. Dynamic exclusion was set to 20 s and peptide match was set to preferred. Mobile phases consisted of 3% acetonitrile/0.1% formic acid as solvent A, 90% acetonitrile/0.1% formic acid as solvent B. Flow rate was set to 200 nL/min throughout the gradient, 2% - 6% B in 1 min, 6% - 30% B in 84 min, 30% - 60% B in 9 min, 60% - 90% B in 1 min with a hold at 90% B for 5 min. MS data were analyzed using Spectrum Mill MS Proteomics Workbench software Rev B.06.01.202 (Agilent Technologies). Similar MS/MS spectra acquired on the same precursor m/z within +/-60 sec were merged. MS/MS spectra were excluded from searching if they failed the quality filter by not having a sequence tag length > 0 (i.e., minimum of two masses separated by the in-chain mass of an amino acid) or did not have a precursor MH+ in the range of 600-6000. All extracted spectra were searched against a UniProt database containing human and mouse reference proteome sequences downloaded from the UniProt web site on October 17, 2014 with redundant sequences removed. A set of common laboratory contaminant proteins (150 sequences) were appended to this database and verified to contain the sequences for human and mouse major prion protein. Search parameters included: ESI-QEXACTIVE-HCD-v2 scoring, parent and fragment mass tolerance of 20 ppm, 40% minimum matched peak intensity and ‘trypsin’ enzyme specificity up to 2 missed cleavages. Fixed modification was carbamidomethylation at cysteine and variable modifications were oxidized methionine, deamidation of asparagine and pyro-glutamic acid. Database matches were autovalidated at the peptide and protein level in a two-step process with identification FDR estimated by target-decoy-based searches using reversed sequences. The list of identified proteins was further filtered to contain proteins and protein isoforms with at least 2 unique peptides and an aggregate protein score greater than 20. Protein-peptide comparison report comprised of all validated peptides was exported which included a ranked summary by intensity of all peptides unique to prion protein.

### Selection of PrP peptides for MRM assay development

Nine peptides covering 4 species were selected from computational and empirical data (Table S2 and Figures S2-S4). Peptides were prioritized based our criteria previously described^33^ and outlined in detail in Figure S2 as well as considerations based on PrP biology and desired assay applications described in Results (Figure 1). One peptide, PIIHFGSDYEDR, was included after being detected in CSF despite an N-terminal proline.

**Figure 1.**
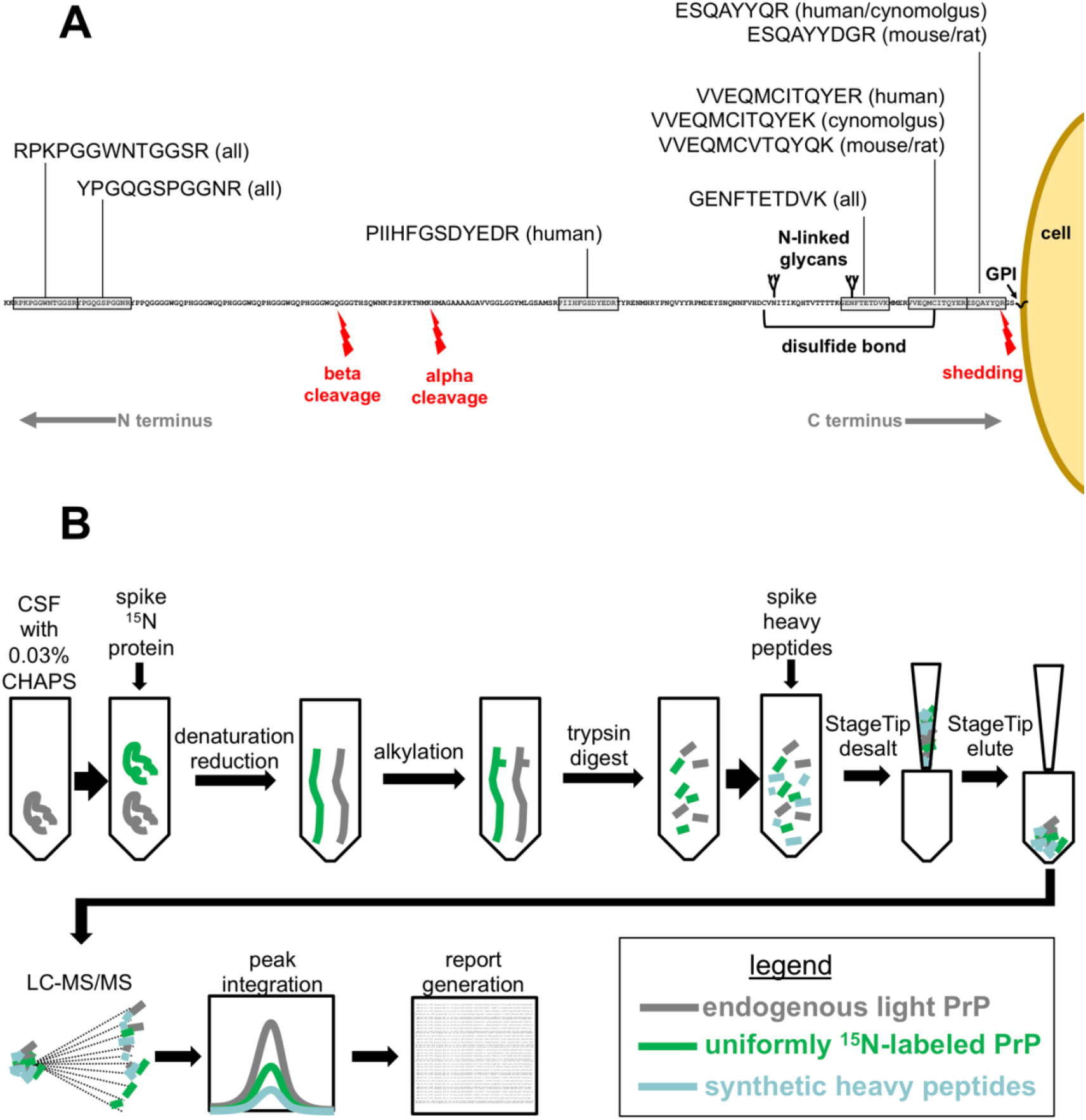
Design of the PrP MRM assay. **A)** Selection of PrP tryptic peptides for MRM. The full sequence of human PrP (residues 23-230) after post-translational modifications (removal of signal peptide residues 1-22 and GPI signal residues 231-253) is shown, GPI-anchored to the outer leaflet of the plasma membrane, with the position of selected peptides and their rodent or monkey orthologs shown relative to the positions of N-linked glycans, a disulfide bond, and endogenous proteolytic events^7^. **B)** PrP MRM workflow as described in Methods.

The nine peptides were synthesized (New England Peptide) using stable isotope labeled [^15^N_4_^13^C_6_]Arg or [^15^N_2_^13^C_6_]Lys at the C terminus and purified peptide specifications previously outlined (>95% chemical purity, >99% isotopic purity, quantified by AAA) in order to qualify as standards for Tier 1 or Tier 2 assays^34^.

### PrP MRM assay

In devising a CSF sample preparation protocol, we drew upon our experience with MRM analysis of plasma^35^ and published mass spectrometry protocols for prion studies^36,37^.

Uniformly labeled ^15^N-labeled recombinant HuPrP23-230 (starting concentration 2.42 mg/mL determined by AAA) with an estimated isotopic incorporation >97.5% (*see Recombinant Protein Preparation*) was diluted 1:5,000 in phosphate-buffered saline containing 1 mg/mL bovine serum albumin and 0.03% CHAPS. This solution was then further diluted 1:20 (1.5 µL added into 30 µL) into CSF samples (final concentration 24.2 ng/mL) prior to the denaturation and digestion workflow described below. ELISA analysis indicated that this concentration of carrier protein and detergent was sufficient to keep recombinant PrP in solution and avoid loss to plastic, without appreciably affecting CSF total protein content.

All concentrations listed below are final concentrations. For each replicate, 30 µL of CSF was incubated with 0.03% CHAPS with 6 M urea (Sigma U0631) and 20 mM TCEP (Pierce 77720) at 37°C while shaking at 800 rpm in an Eppendorf Thermomixer for 30 min to denature the protein and reduce disulfide bonds. 39 mM iodoacetamide was added for 30 min in the dark at room temperature to alkylate cysteine residues. Urea was diluted to 900 mM by the addition of 0.2 Trizma pH 8.1 (Sigma T8568) to permit trypsin activity. 1 µg of trypsin (Promega V5113) was added (final concentration of ∼1.4 ng/µL), providing at least a 1:50 trypsin:substrate ratio for CSF samples with total protein content <1.6 mg/mL, which includes 97% of CSF samples we have analyzed^3^. Trypsin digestion proceeded overnight shaking at 800 rpm at 37°C. Digestion was stopped with 5% formic acid and transfer to 4°C. A mix containing 100 fmol of each ^15^N/^13^C-labeled synthetic heavy peptide was then added to the CSF digests (3.33 nM peptide, equivalent to ∼76 ng/mL full-length PrP based on an approximate molecular weight of 22.8 kDa).

To desalt the samples, StageTips^38^ comprised of two punches of C18 material (Empore 66883-U) fitted into a 200 µL pipette tip using a 16 gauge needle with 90° blunt ends (Cadence Science 7938) and a PEEK tubing puncher (Idex 1567) were placed onto microcentrifuge tubes using an adapter (Glycen CEN.24). Tubes were centrifuged at 2,500*g* for 3 min after each step, as follows: conditioning with 50 µL 90% acetonitrile / 0.1% trifluoroacetic acid; equilibration with 50 µL 0.1% trifluoroacetic acid and priming with 10 µL 0.1% trifluoroacetic acid (no spin after priming); addition of CSF digest in increments of 150 µL; two washes with 50 µL of 0.1% trifluoroacetic acid; and two elutions into a new microcentrifuge tube with 50 µL of 40% acetonitrile / 0.1% trifluoroacetic acid. Eluates were frozen at −80°C.

Frozen samples were dried under vacuum centrifugation and resuspended in 12 µL 3% acetonitrile/5% acetic acid and placed into a vortexer for 5 minutes at room temperature. Samples were then centrifuged at 12,000*g* for 5 minutes and 10 µL of the supernatant was transferred to an HPLC vial (Waters 186000273). HPLC vials were centrifuged briefly (30 - 60s) at 1,200*g* to remove air bubbles and transferred into the nanoLC autosampler compartment set to 7°C. Samples were analyzed on a TSQ Quantiva triple quadrupole mass spectrometer installed with a Nanospray Flex source and Easy-nLC 1000 system (Thermo). Ion source was set to positive ion mode with capillary temperature of 300°C, spray voltage of 2,000 and sweep gas set to 0. The Easy-nLC 1000 system was primed with mobile phase A (3% acetonitrile / 0.1% formic acid), mobile phase B (90% acetonitrile / 0.1% formic acid). Samples were injected (2 µL, 20% of digested sample) onto a 0.075 mm ID PicoFrit (New Objective) column pulled to a 10 µm emitter and custom-packed to 20 cm with 1.9 µm 200Å C18-AQ Reprosil beads (Dr. Maisch). The LC gradient was 0% B to 30% B for 55 min, 30% B to 60% B in 5 min, 60% B to 90 % B in 1 min using a flow rate of 200 nL/min. Collision energies were optimized over 4 steps, 2.5 V per step in batches of less than 500 transitions per batch. Three to four transitions were monitored per peptide using the MRM transitions listed in Table S1 using a 1.5s cycle time. In addition, even though the corresponding heavy peptides were not synthesized, we monitored for the transitions that corresponded to the oxidized methionine version of the peptide VVEQMCITQYER.

### Data analysis

Extracted Ion chromatograms (XIC) of all transition ions were verified and integrated using a Skyline document as described^39^ (Skyline version 4.1.0.11796, https://brendanx-uw1.gs.washington.edu/labkey/project/home/software/Skyline/begin.view) that contained all the selected peptides for the selected species of prion protein. After peak integration, the Skyline report file was exported as a text delimited file where the peak areas in the columns labeled as “Light”, “Heavy” or “15N” for the single most intense, interference-free, reproducibly measured transition (Table S1) were used for quantification and subsequent statistical analysis. Columns included for export were: Protein Name, Protein Gene, Protein Species, Peptide Sequence, Peptide Modified Sequence, File Name, Acquired Time, Replicate Name, SampleGroup, Peptide Retention Time, Precursor Mz, Fragment Ion, Area, Area Ratio, Total Area, Total Area Ratio.

In order to determine the response of each peptide in terms of L:^15^N ratio as well as evaluate dilution linearity of the assay, we spiked 0, 2.4, 24, or 240 ng/mL of ^15^N-labeled recombinant human PrP into a single control CSF sample (from an individual with normal pressure hydrocephalus) in triplicate. For each peptide, we then fitted a linear model correlating the (non-zero) spiked concentrations to the observed ^15^N:light ratios with the intercept fixed at zero, yielding slopes ranging from 39 to 448 ng/mL. Each peptide was then assigned a response factor equal to the highest slope observed for any peptide (448 ng/mL) divided by its own slope.

In N=12 individual replicates (out of 110) of the clinical samples, the oxidized methionine (met-ox) version of the VVEQMCITQYER peptide was more abundant than the reduced version, despite the inclusion of a reduction step in sample preparation. The VVEQMCITQYER peptide was omitted from analysis for these replicates.

Statistical analysis and data visualization were performed using R 3.5.1 in RStudio 1.1.456. Statistical tests are named throughout the text and are all two-sided. Reported *P* values are nominal.

### Data and source code availability

All processed data and source code for this study are provided in a public GitHub repository at https://github.com/ericminikel/prp_mrm and are sufficient to reproduce the analyses and figures herein. This repository also includes a summary table for download containing the MRM results (light and ^15^N peak areas, light:^15^N ratio and normalized PrP concentration in ng/mL) for all clinical samples and all peptides.

## Results

### Design of the PrP MRM assay

PrP ranked number 8 in intensity out of 322 confidently detected proteins in single-shot, LC-MS/MS analysis of human CSF digested with trypsin (see Methods). This indicated that PrP was a good candidate for direct analysis by LC-MRM-MS in CSF without additional fractionation^40^ or enrichment methods^41^. PrP peptides with the highest MS intensities after digestion of recombinant human or mouse PrP as well as human CSF were preferentially ranked according to criteria described in Methods and Figure S2. We selected six human peptides as well as three peptides specific to mouse, rat, and/or cynomolgus macaque PrP, to support assay application to preclinical drug development (Figure 1A, Figure S4, and Table S2). Peptides were chosen to span the N- and C-terminal domains of PrP, up- and down-stream of alpha and beta cleavage sites, allowing us to quantify proteolytic fragments of cleaved PrP (Figure 1A and Figure S4).

We further designed a workflow for the PrP MRM assay (Figure 1B) incorporating an incubation in the presence of a strong chaotrope to denature both properly folded and misfolded forms of PrP. We then reduced and alkylated the protein mixture to break the disulfide bonds and prevent them from refolding, and thereby make the whole protein accessible to the enzymatic processing of r-trypsin. To permit quantification of endogenous, unlabeled (hereafter “light” or “L”) PrP, we added a 9-plex mixture of synthetic ^15^N/^13^C-stable isotope labeled (hereafter “heavy” or “H”) internal standard peptides to CSF samples after digestion. In addition to properly identifying the endogenous light peptides by MRM, these heavy peptides control for variability in retention on the LC and the ionization on the MS, caused by the presence of a large number of peptides in the mixture, with over 4,000 peptides identified in CSF pilot study. To further control for the analytical variability that can occur during enzymatic proteolysis and solid phase extraction (SPE) using StageTips^38^, we also added uniformly ^15^N-labeled recombinant human PrP (hereafter “^15^N”) into clinical samples prior to analysis (Figure 1B).

### Assessment of PrP MRM performance

We conducted a series of analytical validation experiments to assess the performance of the PrP MRM assay. To assess cross-species selectivity and sensitivity, we analyzed human, rat, and cynomolgus macaque CSF as well as mouse and human brain homogenate. For the six PrP peptides harboring sequence differences between species (Figure 1A, Table S2), we observed excellent selectivity, with peptides consistently detected in sequence-matched species above the background level observed in non-sequence-matched species (Figure S5A-B) and with technical replicate mean coefficients of variation (CVs) all <15% (Table S3). In a dose-response experiment, ^15^N-labeled recombinant human PrP added to human CSF in dose-response was recovered with good dilution linearity over at least the two orders of magnitude chosen for this experiment (Figure S5C). Dilution linearity for endogenous CSF PrP was confirmed by mixing high-PrP and low-PrP human CSF samples in different proportions (Figure S5D). We found that the total protein and lipid content of brain tissue precluded analysis of ≥1% brain homogenates, but 0.5% brain homogenates were technically tractable in PrP MRM. Using mixtures of wild-type mouse brain homogenates titrated into a background of PrP knockout mouse brain homogenate, we prepared samples to evaluate the specificity and dilution linearity across a PrP concentration range expected in CSF samples obtained from patients. MRM analysis revealed a linear response for three mouse sequence-matched peptides (Figure S5E).

To support measurement of endogenous unlabeled PrP in *N*=55 human CSF clinical samples (see next section), we performed quality control analysis using the ^15^N protein added into each sample before digestion as well as the cognate synthetic heavy peptides added after digestion. Clinical samples were divided into 5 batches run on separate days; each sample was processed and analyzed in duplicate within its day. A common control sample was also measured in duplicate on each day.

As expected, the mean absolute MS response, either from ^15^N recombinant or from endogenous light PrP, varied by over an order of magnitude between the six PrP peptides (Figure 2A-B), primarily reflecting differences in electrospray ionization efficiencies^40,42,43^. The recovery of the six peptides from endogenous PrP relative to one another was preserved across CSF patient samples (Figure 2A), but differed from the recovery of the corresponding peptides derived from ^15^N recombinant PrP (Figure 2B), resulting in a ∼10-fold difference in mean light:^15^N ratio between different peptides (Figure 2C and Table S5). These differences between peptides were consistent between days (Figure S6), and assessment of the ^15^N:H ratio, which is expected to be the same in all samples, indicated that the analytical process was consistent between samples and days (Table S4). The differences in peptide recoveries may reflect differences in proteolytic processing using our urea/trypsin protocol^44^ and/or post-translational modification (Figure 1A) of PrP in CSF relative to the bacterially expressed recombinant ^15^N version used as reference. For example, a significant proportion of brain PrP is N-terminally truncated^11^, and PrP cleavage products have been observed in CSF as well^10^. PrP is known to be variably glycosylated at residue N197, but our assay will only detect the non-glycosylated form of the GENFTETDVK peptide containing this site. This may account for the much lower response of this peptide in CSF vs. the ^15^N standard (Figure 2). For the C-terminal peptide ESQAYYQR, our assay might not detect proteolytically shed PrP if the cut site for ADAM10, the predominant PrP sheddase^45^, in human PrP is homologous to its reported cut site in rodent PrP^8,46^. For the most N-terminal peptide monitored, RPKPGGWNTGGSR, the presence of a retained N-terminal methionine three residues upstream of this sequence in bacterially expressed PrP, detected here (Figure S3) consistent with reported N-terminal methionine excision patterns in *E. coli*^47^, could alter its trypsin digest efficiency relative to brain and CSF PrP. Because we lacked access to purified full-length mammalian PrP to serve a reference standard, we cannot definitively dissect the reasons for the differences in recovery between peptides. Accordingly, we assigned each peptide a response factor based on the slope of the light:^15^N ratio observed in the ^15^N dose-response experiment (Methods, Figure S8). Applying these response factors to the light:^15^N ratios brought each peptide’s abundance into line with the highest-responding peptide, and yielded estimates of CSF PrP concentration in CSF that averaged 421 ng/mL across samples and all peptides (Table S5, Figure S8).

**Figure 2.**
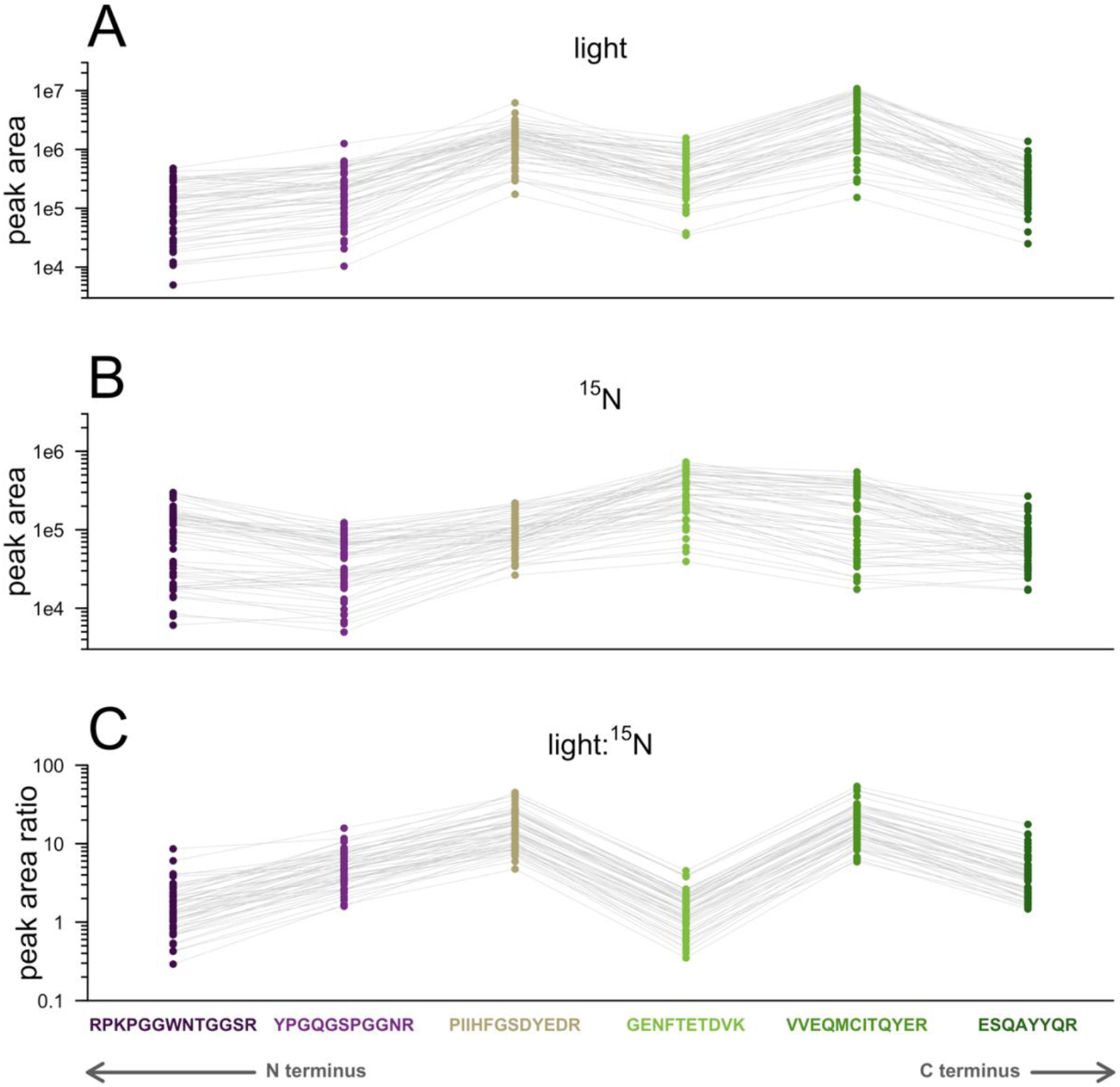
Relative recovery of six human PrP peptides in CSF. For each of N=55 clinical samples, panels show each peptide’s **A)** light peak area, **B)** ^15^N peak area, and **C)** light:^15^N ratio. Gray lines connect the dots representing distinct peptides from the same individual.

All six peptides exhibited strong technical performance on par with other published MRM assays^35,40–42^, with mean same-day technical replicate CVs <15% both overall (Table 1) and within quartiles across the range of low- to high-PrP samples (Table S6), as well as inter-day technical replicate CVs <25%. These data suggest that PrP MRM is suitable for estimating the amount of PrP in CSF and how it changes within and across patients. In further support of the applicability of this multiplex assay to answering biological questions in clinical samples, we found that for every peptide, the variability in amount of PrP between patient samples was much larger than the analytical variability, with inter-individual CVs of 52-80% contrasting with the observed tight technical replicate agreement of ∼10% CV (Table 1). Similar results were obtained when the L:H ratio was used instead (Table S7, Figure S9, S10). Given that analytical variability was much smaller than biological variability, all six peptides were deemed suitable for analysis in clinical samples, and, owing to their different positions within PrP’s amino acid sequence (Figure 1A), each peptide was deemed able to inform independently upon the presence of its particular protein domain in CSF.

**Table 1.**
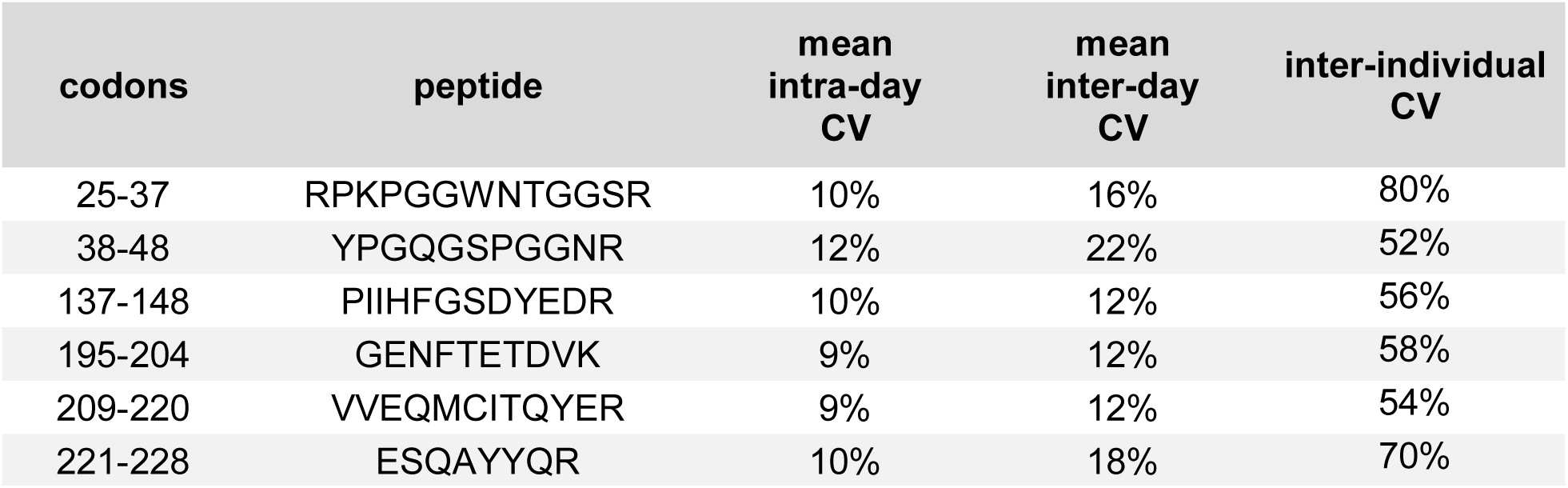
Recovery and performance of six human peptides in human CSF samples. Mean intra-day CV (based on same-day process duplicates of N=55 samples); mean inter-day CV (based on a single inter-day control CSF sample analyzed in duplicate on N=5 separate days; and inter-individual CV among the 55 different samples.

| codons | peptide | mean intra-day CV | mean inter-day CV | inter-individual CV |
| --- | --- | --- | --- | --- |
| 25-37 | RPKPGGWNTGGSR | 10% | 16% | 80% |
| 38-48 | YPGQGSPGGNR | 12% | 22% | 52% |
| 137-148 | PIIHFGSDYEDR | 10% | 12% | 56% |
| 195-204 | GENFTETDVK | 9% | 12% | 58% |
| 209-220 | VVEQMCITQYER | 9% | 12% | 54% |
| 221-228 | ESQAYYQR | 10% | 18% | 70% |

### PrP peptide abundance across diagnostic categories

We used PrP MRM to quantify CSF PrP peptides in *N*=55 clinical samples from individuals with rapidly progressive dementia referred to prion surveillance centers for testing and who ultimately either received non-prion disease diagnoses, or in whom sporadic or genetic prion disease was confirmed by autopsy (see Methods). All six human PrP peptides quantified by PrP MRM showed a marked decrease in abundance in prion disease patients compared to non-prion diagnoses, and all six peptides showed the same general pattern, with non-prion disease patients’ CSF samples giving the highest mean peptide level, followed by sporadic prion disease, followed by genetic prion disease (Figure 3A). The results from MRM mirrored the previously reported PrP ELISA results for these same 55 individuals^3^ (Figure 3B), but differed in the estimated absolute amounts of PrP by ∼3-fold.

**Figure 3.**
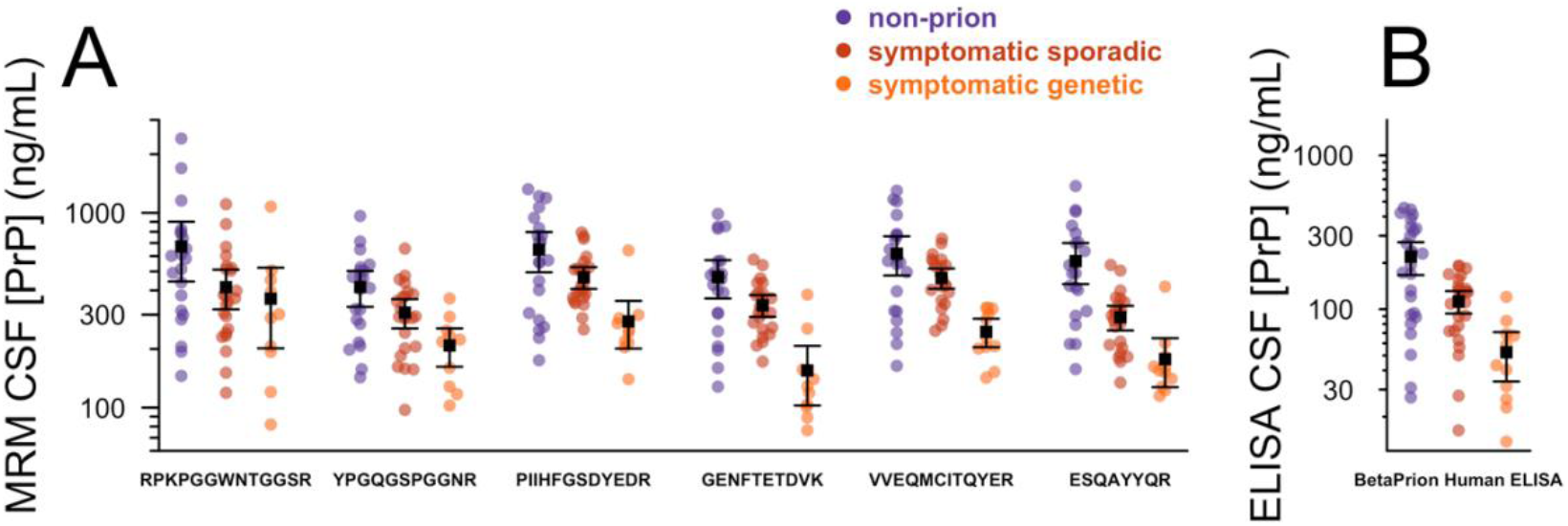
All PrP MRM peptides are decreased in the CSF of prion disease patients. CSF PrP concentrations in N=55 clinical CSF samples determined by **A)** PrP MRM for each of six peptides, arranged with the most N-terminal peptide at left and the most C-terminal peptide at right, compared with **B)** previously reported PrP ELISA results for the same samples, reproduced from Vallabh et al^3^. Black squares and bars show the mean and 95% confidence interval of the mean for each group.

### Relationship between PrP MRM and ELISA

Across the clinical samples, each peptide’s abundance was positively correlated to the full-length PrP concentration determined by ELISA (Figure 4A). The coefficients of correlation, from 0.40 to 0.72, are within the ranges reported for other MRM assays compared to corresponding immunoassays^41,42,48^. All peptides were strongly correlated to one another, with coefficients of correlation ranging from 0.67 to 0.96, and no obvious differences within versus between protein domains (N- and C-terminal; Figure 4B). The linear relationships between peptides were preserved across the range of samples analyzed and were similar in terms of both L:H as well as L:^15^N ratios (Figure S10). These results, together with the fact that the magnitude of decrease in abundance in prion disease cases was similar for all peptides (Figure 4A), suggested that PrP MRM and ELISA may be measuring the same analyte — predominantly full-length PrP. We therefore asked whether PrP MRM could serve as an orthogonal method to validate findings recently reported for ELISA.

**Figure 4.**
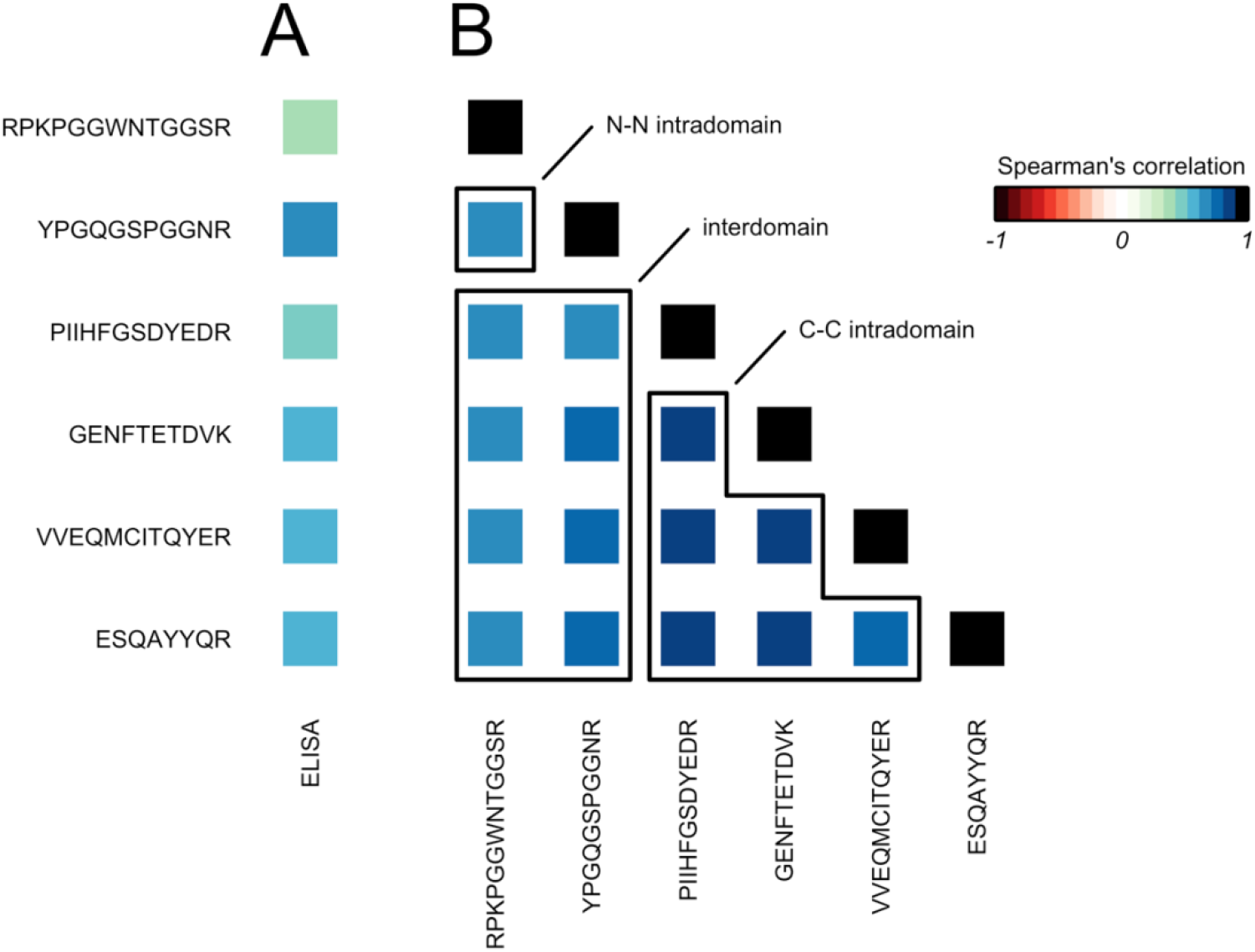
Correlations among PrP MRM peptides and with ELISA. **A)** Spearman’s correlation between each peptide measured in MRM versus total PrP by ELISA. **B)** Spearman’s correlation between every combination of peptides measured in MRM. All P < 0.01.

Because plastic adsorption is reported to cause substantial loss of PrP in preanalytical handling, and detergent is reported to largely mitigate this^3^, we analyzed replicates of one CSF sample by MRM with and without 0.03% CHAPS detergent. As with ELISA, we found that the addition of CHAPS increased PrP peptide recovery by an average of 51% (*P* = 2.3e-8, Type I ANOVA).

To compare PrP MRM and ELISA results while introducing covariates, we calculated a final estimated PrP concentration from MRM for each CSF sample by averaging the normalized PrP concentration across the six peptides. The estimated PrP concentrations obtained by MRM and by ELISA were correlated across CSF samples (r = 0.61, Spearman’s correlation, *P* = 1.3e-6). MRM PrP concentration was uncorrelated with CSF hemoglobin (*P* = 0.85, Spearman’s correlation), supporting the conclusion that blood contamination is not a source of CSF PrP^3^.

The concentration of PrP in CSF measured by ELISA is correlated with the total protein concentration in CSF^3^. This could reflect true biology, or it could reflect pre-analytical factors, if other proteins serve a blocking function, mitigating PrP loss to plastic during handling^3^. A potential concern, however, is that such a correlation could also arise if non-specific binding of other proteins in the human CSF matrix contributes to PrP ELISA background signal. If true, this would call into question the ability of ELISA-based PrP measurement to accurately quantify a pharmacodynamic decrease in PrP concentration. To distinguish between these possibilities, we tested the relationships between ELISA PrP concentration, MRM PrP concentration, and total protein concentration among our clinical samples. The correlation between ELISA PrP concentration and total protein concentration was marginal but observable among the 55 samples analyzed here (+94 ng/mL PrP per 1 mg total protein, *P* = 0.043, linear regression: ELISA PrP ∼ total protein), but this relationship vanished completely when MRM PrP concentration was included as a covariate (*P* = 0.60 for total protein in linear regression: ELISA PrP ∼ MRM PrP + total protein). Likewise, MRM PrP concentration was itself correlated to total protein (+238 ng/mL PrP per 1 mg/mL total protein, *P* = 0.017, linear regression: MRM PrP ∼ total protein). Together, the observations that the relationship between PrP and total protein was replicated in MRM, and that total protein did not explain any residual variance in ELISA-measured PrP after controlling for MRM-measured PrP, suggest that the correlation between CSF PrP and total protein in CSF is a genuine property of the samples analyzed, and that ELISA is specifically measuring PrP in human CSF.

## Discussion

Here we describe a targeted mass spectrometry assay for measuring CSF PrP. Six of six human PrP peptides we quantified, from the N to the C terminus, were lowered in prion disease patients compared to non-prion disease patients. Thus, the highly reproducible finding that CSF PrP concentration decreases in prion disease^3,13–16^ appears to represent genuine disease biology and is not merely an ELISA measurement artifact. This confirms that CSF PrP will be difficult to interpret as a pharmacodynamic biomarker in symptomatic prion disease patients, because the direct effect of a PrP-lowering drug and the effect of disease process alleviation would be expected to push CSF PrP in opposing directions. Instead, trials to demonstrate target engagement and perform dose-finding for a PrP-lowering drug may need to be conducted in pre-symptomatic individuals at risk for genetic prion disease^2,3^.

We also validate other findings from PrP ELISA studies. We confirm that the correlation between CSF PrP and total protein is genuine, and not just a result of matrix interference in ELISA. We also confirm that CSF PrP is not correlated with CSF hemoglobin, further supporting the brain and not blood origin of CSF PrP. Our data provide supportive evidence for the existing literature indicating that CSF PrP can be meaningfully quantified by ELISA.

Our study has several limitations. First, we have only compared samples between prion and non-prion disease patients to examine the effect of the disease state on CSF PrP. Determining the effect of PrP-lowering drug treatment on CSF PrP is a priority for future work. Second, we still cannot exclude the possibility that protein misfolding contributes somewhat to the decrease in CSF PrP that we observe, because the chaotrope used here — 6 M urea — has not been proven to denature all misfolded PrP. This concentration of urea was shown to abolish 99.99% of hamster prion infectivity^49^, but prion strains differ in their conformational stability^24^. Human prions unfold at ∼3 M guanidine hydrochloride^50–52^, but urea is a less potent denaturant^53^. Third, while our assay appears to perform very well, we have not undertaken the full bioanalytical method validation that would be expected if the assay is to be used in clinical decision-making^54^, and the LC/MS gradient used here, at 45 minutes, is longer than the ∼5 minutes expected for high-throughput clinical biomarker assay. For clinical use, the feasibility and performance of the assay would likely need to be assessed at a faster gradient under microflow conditions using commercially available C18 columns. This increase in assay throughput may come at the cost of some sensitivity, but because all PrP peptides in this study demonstrated comparable behavior across this set of clinical samples, a future implementation of PrP MRM might choose to monitor fewer or even a single peptide, facilitating the implementation of a chromatographically faster procedure. Fourth, because bacterially expressed recombinant PrP is an imperfect standard by which to quantify mammalian PrP, our data do not support any firm conclusions about the baseline composition of PrP in terms of different cleavage products in human CSF generally. Nevertheless, by comparing the abundance of each PrP peptide between diagnostic categories — individuals with and without prion disease — we do establish that any changes driven by the disease state apparently affect all domains of PrP equally. This finding is not inconsistent with existing literature: for example, the PrP C2 fragment resulting from beta cleavage is known to be increased in brain parenchyma during prion disease^25^, but if C2 is then retained in intracellular aggregates rather than being shed, while its counterpart N2 is rapidly degraded, then increased beta cleavage might result in both N- and C-terminal PrP peptides being decreased in prion disease CSF, as observed here. Our principal finding, that all PrP peptides move in concert with one another in the disease state, contrasts with the more complex situation reported for tau isoforms in CSF^55,56^, and should simplify the use of CSF PrP quantification as a tool in drug development.

As PrP-lowering therapies progress towards the clinic, MRM and ELISA both appear suitable as tools for measuring PrP in CSF. ELISA is cheaper and less equipment-intensive. MRM offers a wide dynamic range without dilution, and applicability of a single assay both to humans and to multiple preclinical species of interest. Regardless of whether MRM or ELISA is ultimately used in preclinical development and clinical testing of PrP-lowering drugs, the concordance between the two methods builds confidence in CSF PrP as an analyte, and supports its use as a pharmacodynamic biomarker and, perhaps, as a trial endpoint.

## Acknowledgments

This study was supported by the Next Generation Fund at the Broad Institute, BroadIgnite, the National Institutes of Health (F31 AI122592 to EVM), the National Science Foundation (GRFP 2015214731 to SMV), individual philanthropic donations to the Broad Institute and to Prion Alliance, and an anonymous organization. Sample collection in Göttingen was supported by Bundesministerium für Gesundheit (grant 1369-341) through Robert Koch Institute.

## Author contributions

Conceived and designed the study: EVM, SMV, EK, SLS, SAC

Performed the experiments: EVM, EK, ARC, SMV, CRH, AGR

Collected samples and/or clinical data: FL, IZ, SC, PP, GJR, JGS

Supervised the research: SLS, SAC, ROK, MDM

Wrote the paper: EVM, EK

Contributed to experimental design and/or interpretation: all authors

Reviewed and edited the paper: all authors

## Supplementary Materials

**Table S1.**
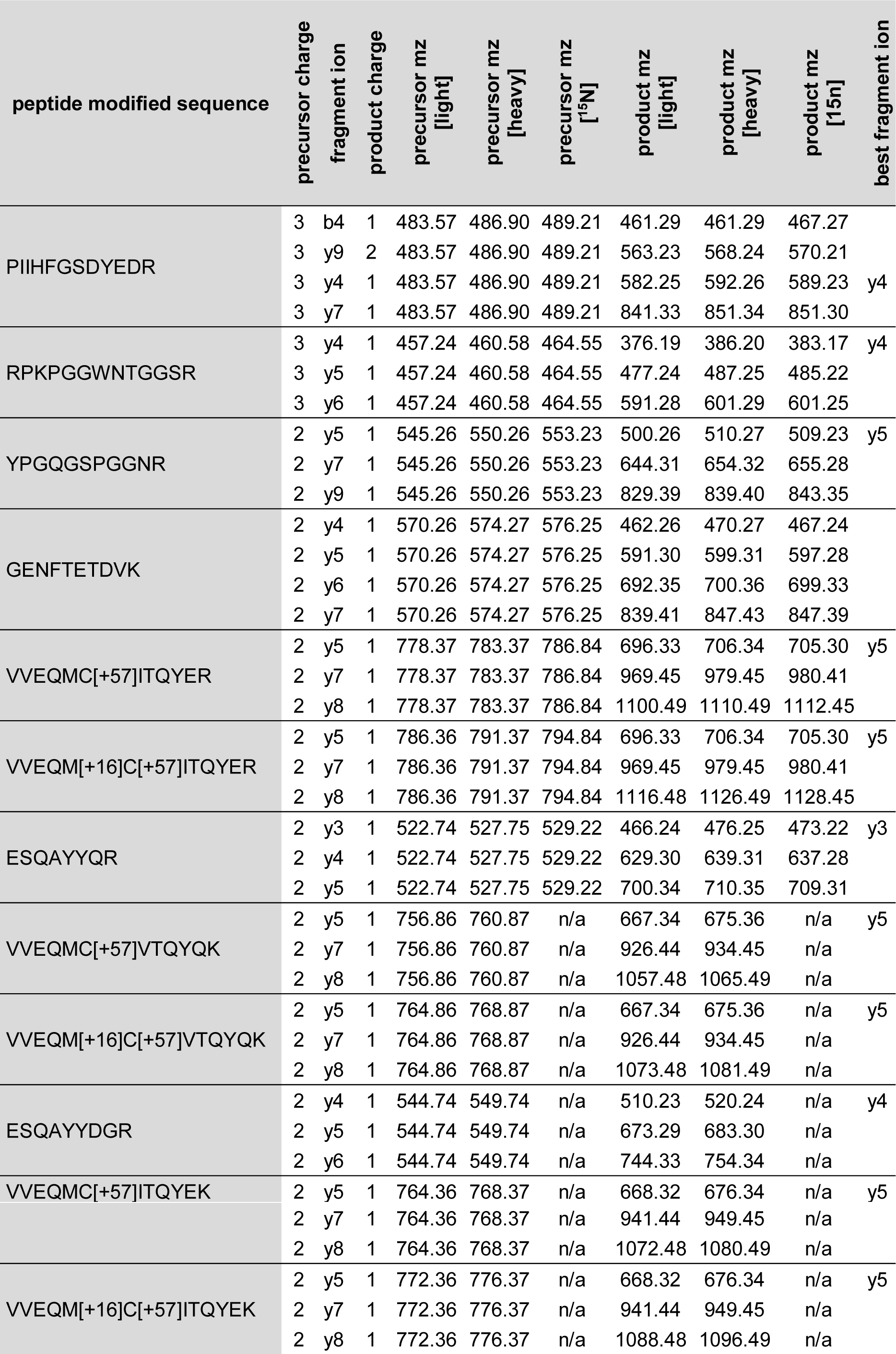
Precursor and product characteristics for all peptides monitored. Best fragment ions were chosen as transition ions with the highest peak area that were also interference-free and reproducibly measured in pilot studies.

**Table S2.** Species sequence matching, sequence context, and pilot study detection of all peptides monitored.

| previous AA | sequence | next AA | species | MS peak area in recombinant PrP | peak area in CSF |
| --- | --- | --- | --- | --- | --- |
| (LCKK) | RPKPGGWNTGGSR | (YPGQ) | human, cyno, mouse, rat | 5.49E+09 | 1.24E+08 |
| (GGSR) | YPGQGSPGGNR | (YPP) | human, cyno, mouse, rat | 4.90E+10 | 8.62E+08 |
| (AMSR) | PIIHFGSDYEDR | (YYR) | human |  | 1.06E+09 |
| (TTTK) | GENFTETDVK | (MME) | human, cyno, mouse, rat | 5.11E+10 | 2.48E+08 |
| (MMER) | VVEQMCITQYER | (ESQ) | human | 3.46E+09 | 1.12E+09 |
| (MMER) | VVEQMCVTQYQK | (ESQA) | mouse, rat | 5.21E+10 |  |
| (MMER) | VVEQMCITQYEK | (ESQ) | cyno |  |  |
| (QYER) | ESQAYYQR | (GSS) | human, cyno | 2.32E+10 |  |
| (QYQK) | ESQAYYDGR | (RSS) | mouse, rat | 6.85E+10 |  |

**Table S3.** Characteristics and performance of the nine PrP MRM peptides in assay development samples. Analytical validation experiments were performed without ^15^N protein internal controls and instead utilized the light:heavy peptide area under the curve ratio as described in Methods. Data from N=19 samples (N=4 cynomolgous macaque CSF, N=10 human CSF, N=1 human brain, N=1 mouse brain, and N=4 rat CSF) in a total of N=35 replicates were analyzed to determine the basic performance characteristics of each peptide (this table) as well as the sensitivity and selectivity of the assay (Figure S5). Here, only data from samples where the peptide is sequence-matched to the species in question are shown. *Reduced peptides only; met-ox versions were not monitored in these runs. Retention time is shown as mean±sd in minutes for a 45-minute gradient. Mean L:H ratio is the mean light:heavy area ratio. Mean CV is calculated across the subset of samples run in technical duplicate or triplicate within the same run, and N indicates the number of unique samples.

| peptide | retention time (min) | mean L:H ratio | mean CV |
| --- | --- | --- | --- |
| RPKPGGWNTGGSR | 16.9±0.7 (N=28) | 0.1 (N=21) | 11% (N=12) |
| YPGQGSPGGNR | 18.6±0.7 (N=28) | 0.6 (N=21) | 5.2% (N=12) |
| PIIHFGSDYEDR | 36.5±1.8 (N=19) | 3.9 (N=13) | 6.5% (N=9) |
| GENFTETDVK | 26.7±1.3 (N=28) | 0.4 (N=21) | 7.2% (N=12) |
| VVEQMCITQYER* | 37.0±1.5 (N=19) | 7.0 (N=13) | 5.7% (N=9) |
| VVEQMCVTQYQK* | 33.7±0.6 (N=3) | 1.0 (N=4) | 7.2% (N=1) |
| VVEQMCITQYEK* | 35.3±0.3 (N=6) | 1.4 (N=2) | 4.3% (N=2) |
| ESQAYYQR | 19.9±0.8 (N=25) | 2.2 (N=17) | 3.3% (N=11) |
| ESQAYYDGR | 23.3±0.3 (N=3) | 0.3 (N=4) | 0.92% (N=1) |

**Table S4.** Analytical process variability assessment. Because the spiked ^15^N protein and synthetic heavy peptide concentrations were the same for every sample and every day, we used the ^15^N:H peak area ratio (PAR) to assess analytical process variability. Mean PARs varied by no more than 22% per day (largest variability: VVEQMCITQYER, 0.28 for day 4 vs. 0.23 for day 3) and mean CVs among all samples, all replicates within each day was <20% for all days and <15% for all days except YPGQGSPGGNR day 3. These data provide supporting evidence for the technical validity of our analytical process, and suggest that the assay is suitable to measure the biological variability among our clinical samples. *Excludes N=12 met-ox samples.

| peptide | <sup>15</sup>N:H PAR mean CV by day |  |  |  |  | <sup>15</sup>N:H mean PAR by day |  |  |  |  |
| --- | --- | --- | --- | --- | --- | --- | --- | --- | --- | --- |
|  | 1 | 2 | 3 | 4 | 5 | 1 | 2 | 3 | 4 | 5 |
| RPKPGGWNTGGSR | 5.7% | 8.2% | 11.0% | 10.4% | 6.9% | 0.04 | 0.05 | 0.04 | 0.03 | 0.03 |
| YPGQGSPGGNR | 8.0% | 13.9% | 18.4% | 13.5% | 9.4% | 0.08 | 0.08 | 0.08 | 0.07 | 0.08 |
| PIIHFGSDYEDR | 13.5% | 10.6% | 10.7% | 9.1% | 12.5% | 0.13 | 0.13 | 0.13 | 0.13 | 0.14 |
| GENFTETDVK | 7.4% | 9.4% | 9.5% | 4.5% | 8.2% | 0.24 | 0.23 | 0.24 | 0.21 | 0.21 |
| VVEQMCITQYER* | 6.9% | 8.2% | 13.9% | 7.8% | 9.4% | 0.26 | 0.25 | 0.23 | 0.28 | 0.25 |
| ESQAYYQR | 7.4% | 5.7% | 12.1% | 10.5% | 9.0% | 0.30 | 0.28 | 0.31 | 0.32 | 0.25 |

**Table S5.** Normalization of peptide responses. The L, ^15^N, and L:^15^N columns summarize the data from Figure 2. Across N=55 clinical samples, the mean L:^15^N ratio for each peptide varied by >10-fold. If these ratios are simply multiplied by the known concentration of ^15^N protein spiked in (24 ng/mL), they correspond to “raw” PrP concentrations here, ranging from 39 – 478 ng/mL. We calculated a response factor for each peptide as described in Methods and Figure S8, which, for the one CSF sample used in the dose-response experiment, serves to bring each peptide up to equal abundance as the highest-responding peptide (VVEQMCITQYER). Multiplying the L:^15^N ratio by the response factor and the spiked ^15^N PrP concentration in clinical samples yields normalized PrP concentrations that are within ±50% of one another. Note that this small residual difference between peptides in terms of normalized PrP concentration reflects the fact that the single CSF sample used in the dose-response experiment did not have exactly the same ratio among the different peptides as the average clinical sample.

| peptide | mean ± sd<br>L peak<br>area<br>(millions) | mean ± sd<br><sup>15</sup>N peak<br>area<br>(millions) | mean ± sd<br>L:<sup>15</sup>N ratio | mean<br>raw<br>[PrP]<br>(ng/mL) | response<br>factor | mean<br>normalized<br>[PrP]<br>(ng/mL) |
| --- | --- | --- | --- | --- | --- | --- |
| RPKPGGWNTGGSR | 0.14±0.12 | 0.10±0.08 | 1.8±1.4 | 43 | 11.6 | 503 |
| YPGQGSPGGNR | 0.26±0.22 | 0.05±0.03 | 5.3±2.7 | 130 | 2.5 | 329 |
| PIIHFGSDYEDR | 1.59±1.04 | 0.10±0.05 | 16.7±9.3 | 409 | 1.2 | 497 |
| GENFTETDVK | 0.49±0.36 | 0.33±0.19 | 1.6±0.9 | 39 | 9.0 | 350 |
| VVEQMCITQYER | 3.67±3.17 | 0.20±0.15 | 19.7±10.7 | 478 | 1.0 | 478 |
| ESQAYYQR | 0.35±0.28 | 0.08±0.05 | 4.8±3.3 | 116 | 3.2 | 373 |

**Table S6.** Characteristics and performance of the six PrP MRM human peptides in clinical samples by quartile. For each peptide, the N=55 samples were broken into quartiles of L:^15^N ratio. The mean coefficient of variation (CV) of technical duplicates and the mean L:^15^N ratio of samples was calculated within each quartile. Note that the rank order used for binning is similar (Figure 3B) but not identical between peptides. Also note that for VVEQMCITQYER, because 12 replicates (including both replicates of one sample) with methionine oxidation were thrown out, sample size is N=44 for CV calculations (using only those samples with N=2 valid process replicates) and N=54 for mean L:^15^N ratio calculations (including all samples with N≥1 valid process replicate). The results show that all six peptides, across all four quartiles, had mean CV ≤15% and mean L:^15^N ratio ≥0.6. Because our data suggest assay linearity extending at least as low as 0.1x of the ^15^N PrP concentration we used (estimated to be 0.24 ng/mL, Figure S5C), this suggests that PrP MRM had acceptable performance in all quartiles and that all measurements in clinical samples were within the dynamic range of the assay.

| peptide | mean CV by quartile |  |  |  | mean L:<sup>15</sup>N ratio by quartile |  |  |  |
| --- | --- | --- | --- | --- | --- | --- | --- | --- |
|  | 0-24% | 25-49% | 50-74% | 75-100% | 0-24% | 25-49% | 50-74% | 75-100% |
| RPKPGGWNTGGSR | 9.8% | 14.5% | 11.7% | 3.9% | 0.7 | 1.2 | 1.8 | 3.7 |
| YPGQGSPGGNR | 12.5% | 11.9% | 14.0% | 9.5% | 2.5 | 4.1 | 5.9 | 9.3 |
| PIIHFGSDYEDR | 10.9% | 11.3% | 5.4% | 11.1% | 7.9 | 11.9 | 17.8 | 30.9 |
| GENFTETDVK | 12.1% | 9.9% | 8.8% | 6.6% | 0.6 | 1.2 | 1.8 | 2.9 |
| VVEQMCITQYER | 12.8% | 12.0% | 3.4% | 6.4% | 9.0 | 14.9 | 21.7 | 35.6 |
| ESQAYYQR | 9.2% | 9.3% | 10.7% | 9.1% | 2.0 | 3.1 | 4.8 | 9.6 |

**Table S7.** Recovery and performance of six human peptides quantified in human CSF samples using L:H ratio. This table is identical to Table 1 except using the L:H peak area ratio rather than the L:^15^N peak area ratio.

| codons | peptide | mean intra-run CV | mean inter-run CV | inter-individual CV |
| --- | --- | --- | --- | --- |
| 25-37 | RPKPGGWNTGGSR | 6% | 25% | 63% |
| 38-48 | YPGQGSPGGNR | 7% | 13% | 55% |
| 137-148 | PIIHFGSDYEDR | 7% | 8% | 59% |
| 195-204 | GENFTETDVK | 5% | 11% | 57% |
| 209-220 | VVEQMCITQYER | 5% | 9% | 58% |
| 221-228 | ESQAYYQR | 4% | 6% | 66% |

**Figure S1.**
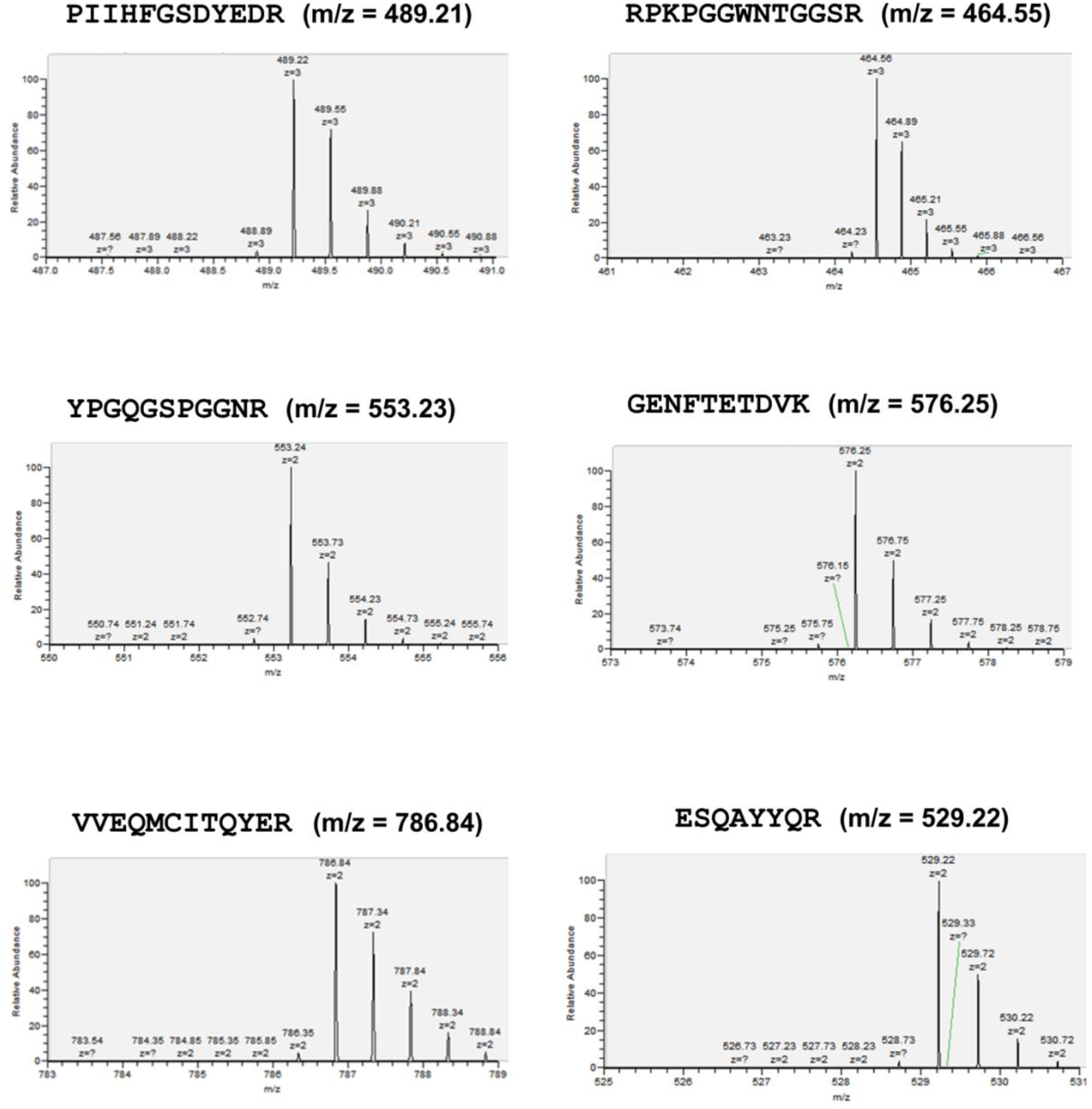
Extracted MS intensities for human PrP peptides used for estimation of isotopic purity of ^15^N-labeled protein. Isotopic envelopes of each peptide identified by MS/MS after digestion of the ^15^N protein with trypsin. Minimal or lack of observed mz peak areas less than the ^12^C monoisotopic mass peak (highest signal for the mz of these peptides) indicates near complete ^15^N incorporation. Lower mass peaks corresponding to incomplete ^15^N incorporation were unquantifiably small, consistent with >97.5% isotopic purity for all peptides.

**Figure S2.**
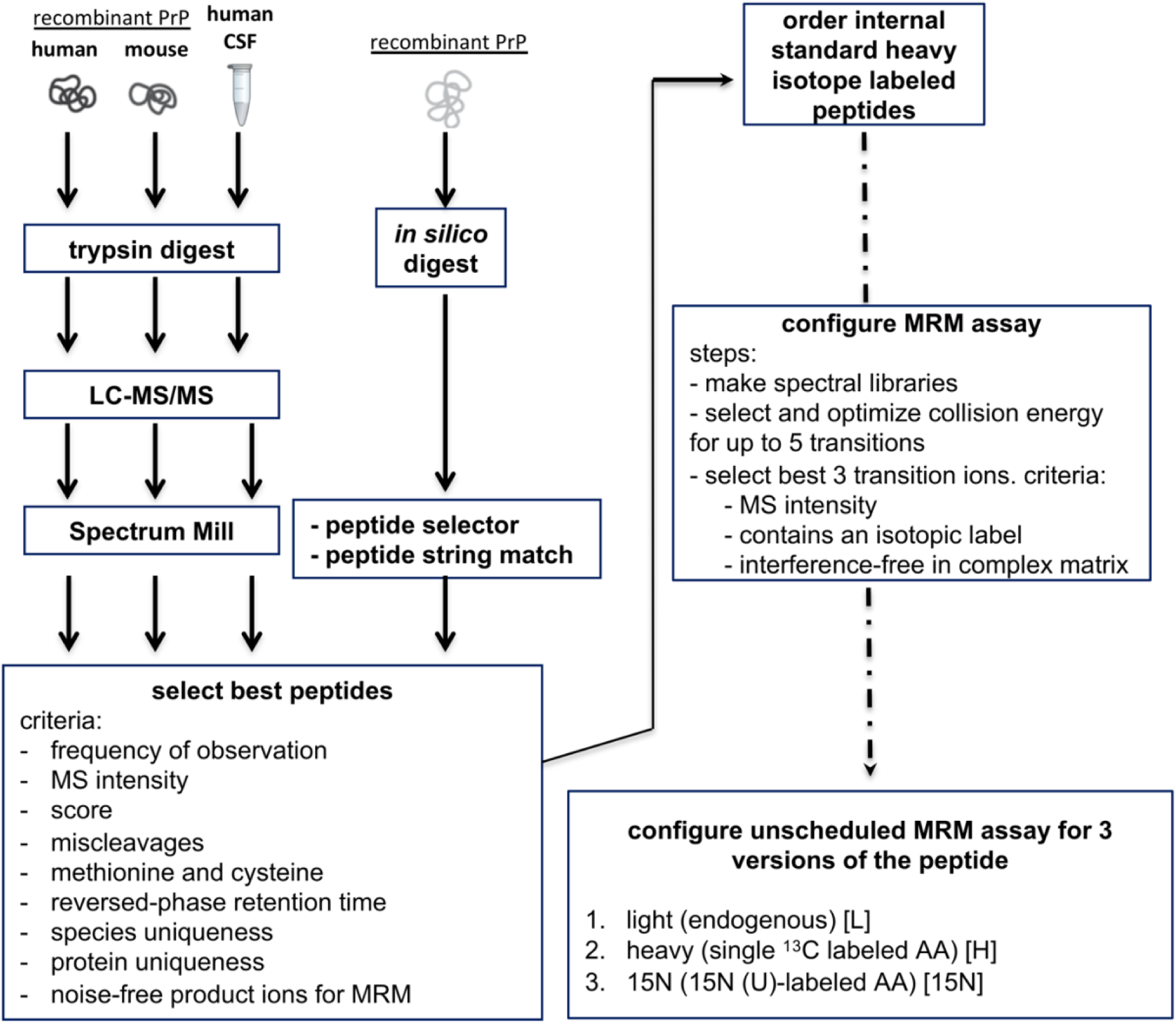
Assay development workflow. Schematic outline of steps described in Methods to select peptides based on empirical and bioinformatic data, and optimize and configure a 9-plex MRM assay.

**Figure S3.**
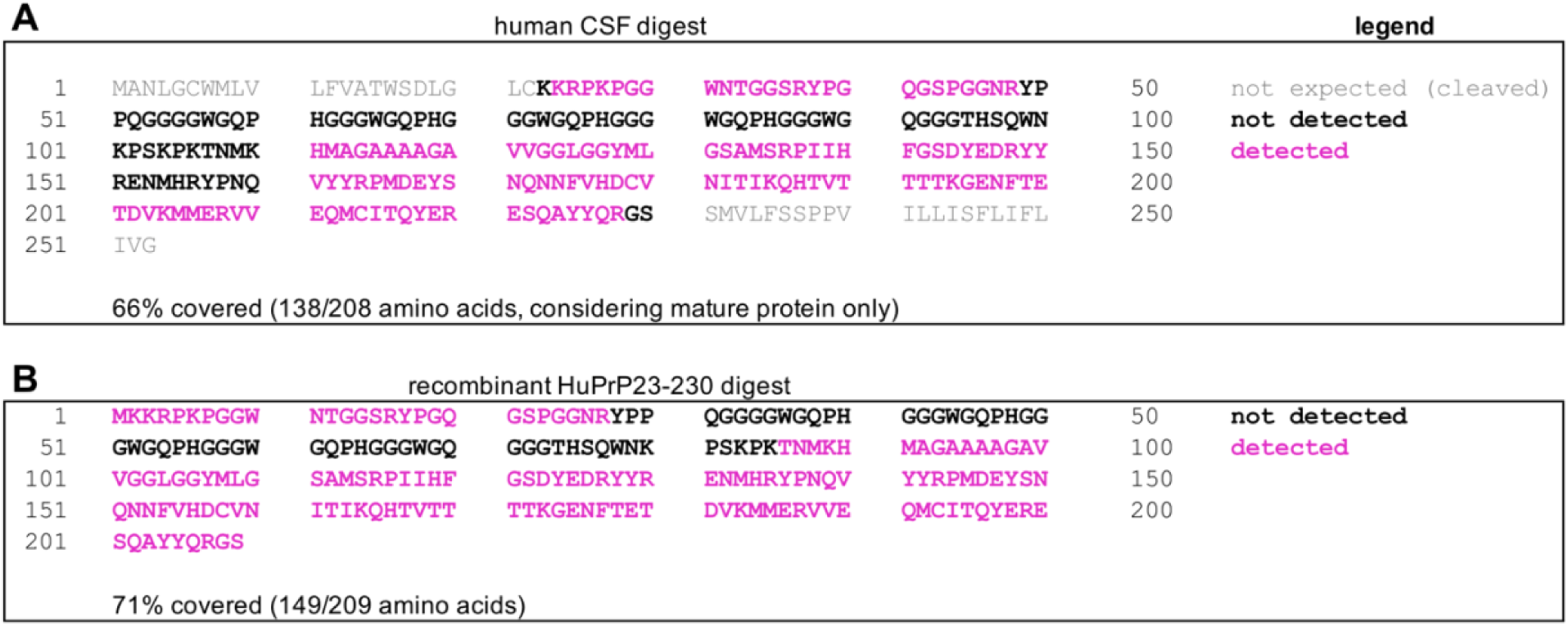
Sequence coverage map. Map of sequence coverage of PrP in pilot LC-MS/MS analyses of **A)** human CSF and **B)** recombinant HuPrP23-230. A peptide containing a retained N-terminal methionine (MKKRPKPGGWNTGGSR) was detected in the recombinant digest with intensity 6.9×10^9^.

**Figure S4.**
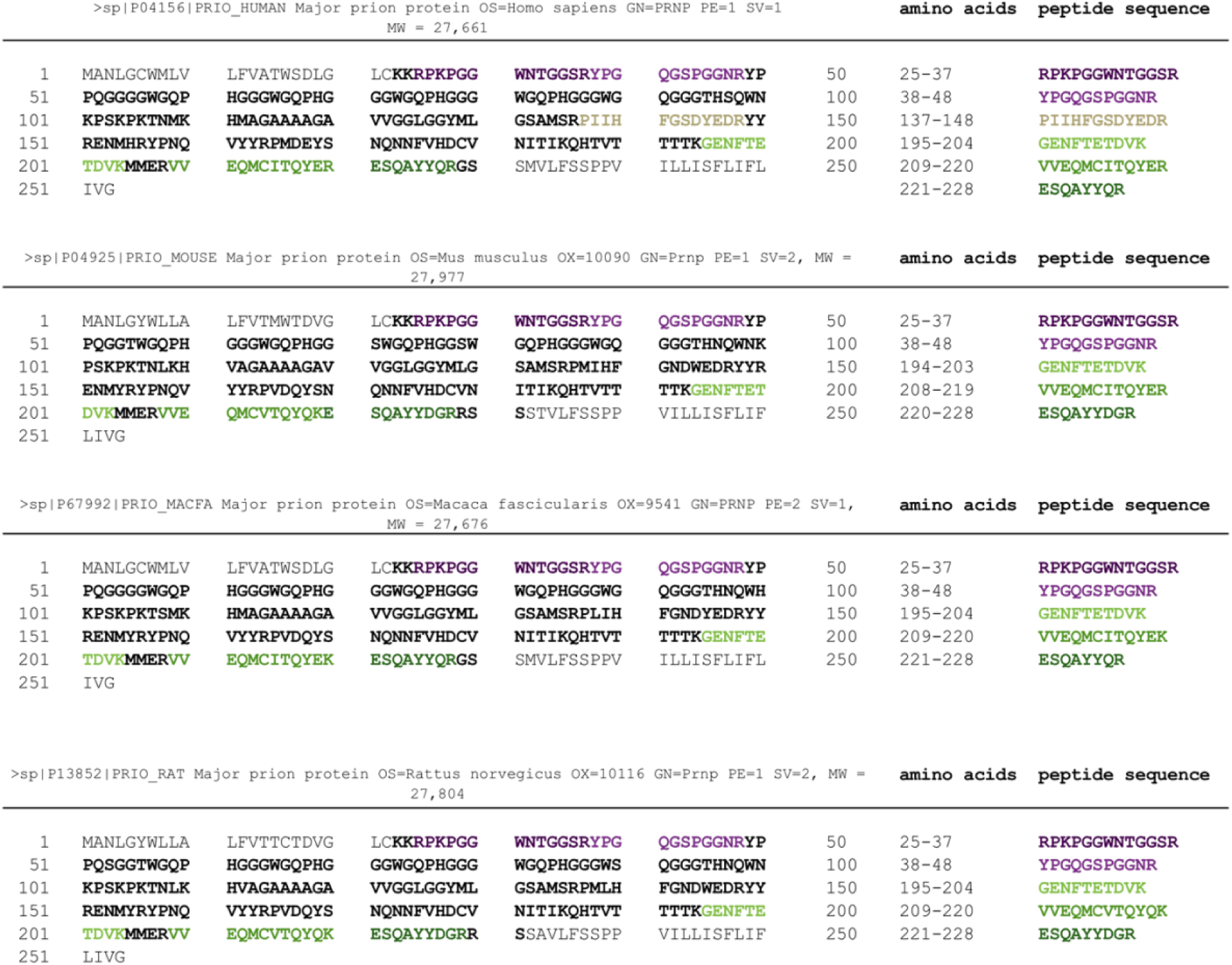
Selected PrP peptides in protein context. Full amino acid sequences of PrP with UniProt identifiers for the four species of interest with locations of PrP MRM peptides noted. Bold indicates residues present in the mature, post-translationally modified protein. The non-bold N terminus is an ER signal peptide, and the non-bold C terminus is a GPI signal, and both are cleaved before the protein reaches the cell surface. Molecular weights pulled from UniProt do not account for these post-translational modifications; the mature protein is ∼23 kDa, see Methods in main text. For human and mouse PrP, the bold text also corresponds to the residues present in the recombinant PrP constructs used in this study.

**Figure S5.**
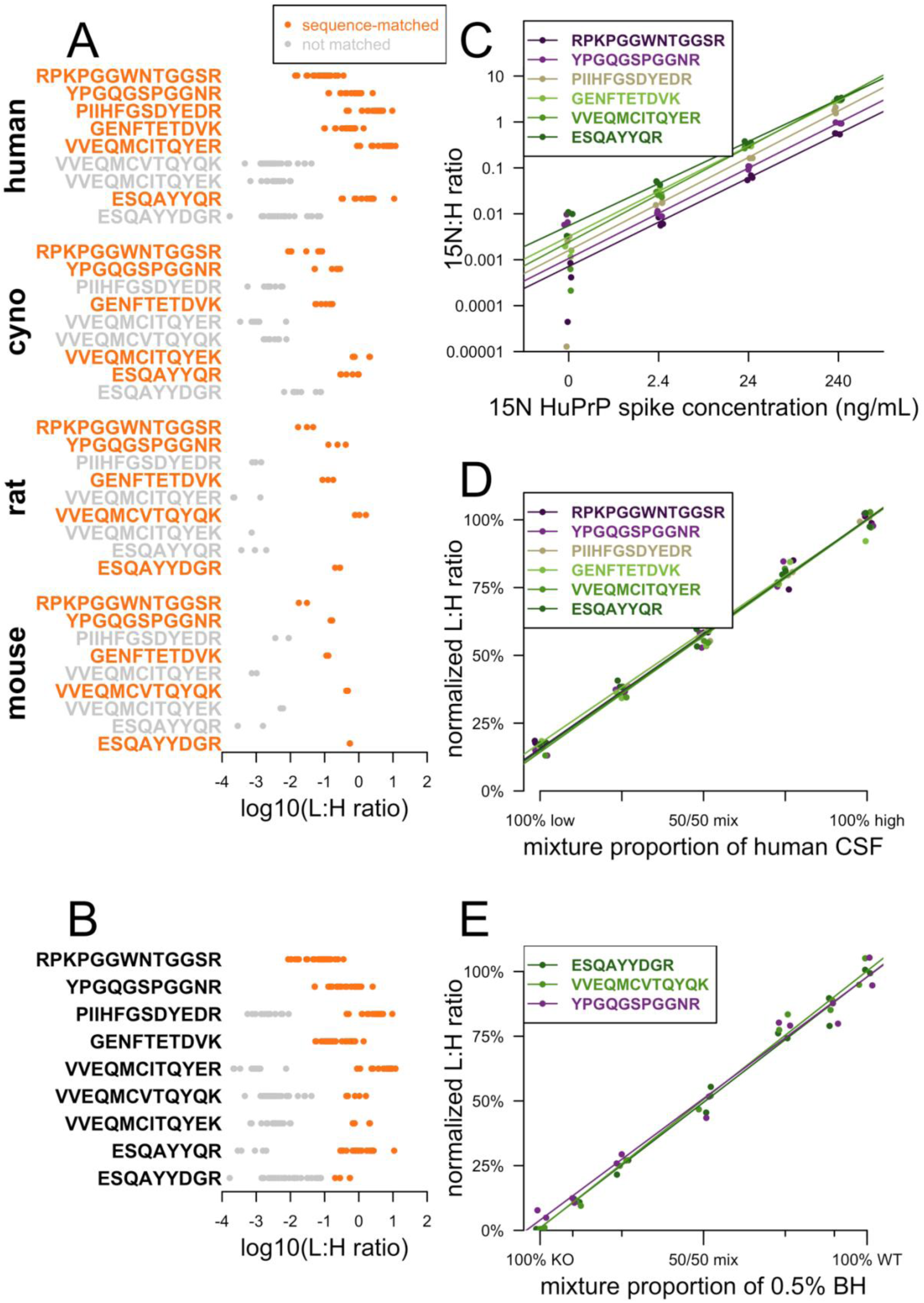
Partial analytical validation of the PrP MRM assay. **A)** Sensitivity and selectivity across species. Data from N=19 samples (N=4 cynomolgous macaque CSF, N=10 human CSF, N=1 human brain, N=1 mouse brain, and N=4 rat CSF) in a total of N=35 replicates were analyzed. L:H peptide ratios are shown for peptides expected in each species (sequence-matched, orange) versus not expected (non-matched, gray). **B)** Results from panel A collapsed across species. This shows that all species-specific peptides were observed in the sequence-matched species at least an order of magnitude above the noise observed in non-sequence-matched species, with the exception of ESQAYYDGR (sequence-matched species: mouse, rat), for which the separation was only about half an order of magnitude. **C)** Assay linearity. ^15^N HuPrP23-230 was spiked into the same human CSF sample in duplicate at three concentrations plus a zero (x axis) and quantified relative to heavy (H) peptides. Best fit lines for each peptide are shown. Peptides vary in absolute recovery (different y-intercepts), as also shown in Table 1, but exhibit similar strong linearity (slopes range 0.96 – 1.04 and adjusted R^2^ values range 99.5% - 99.8%, linear regression), suggesting that normalization relative to ^15^N internal control should provide at least 2 orders of magnitude dynamic range for endogenous PrP. The ^15^N:L ratio in this experiment was used to calculate response factors for each peptide, see Methods. **D)** Two human CSF samples previously measured to have high (240 ng/mL) and low (12 ng/mL) PrP by ELISA were mixed in different proportions (all low, 25/75, 50/50, 75/25, and all high) and assayed by PrP MRM. Each peptide’s light:heavy ratio is normalized to the average value of the two “all high” replicates, and best-fit lines are shown. Individual replicates are jittered slightly along the x-axis so that separate points are visible. Each peptide exhibits good linearity. Note that because the low-PrP CSF sample still has non-zero PrP, the fact that the y-intercepts are non-zero is expected. Best fit lines for each peptide have adjusted R^2^ values ranging 97.6% - 99.8% (linear regression). **E)** 10% brain homogenate from wild-type mice (WT) or Edinburgh PrP knockout mice^29^ (KO) were mixed in seven different proportions (all KO, 10/90, 25/75, 50/50, 75/25, 90/10, and all WT), further diluted to 0.5% brain homogenate in saline and 0.03% CHAPS, and assayed by PrP MRM. Of the five peptides sequence-matched to mouse PrP, the three with best performance in this experiment (mean process replicate CV <10%) are shown here, again with individual replicates jittered along the x axis so that separate points are visible. Each peptide’s L:H ratio is normalized to the average value of the two “all WT” replicates, and best-fit lines are shown. All three peptides exhibit good linearity, with y-intercepts very close to zero, as expected for PrP knockout mice, and adjusted R^2^ values ranging 98.2% - 99.0% (linear regression).

**Figure S6.**
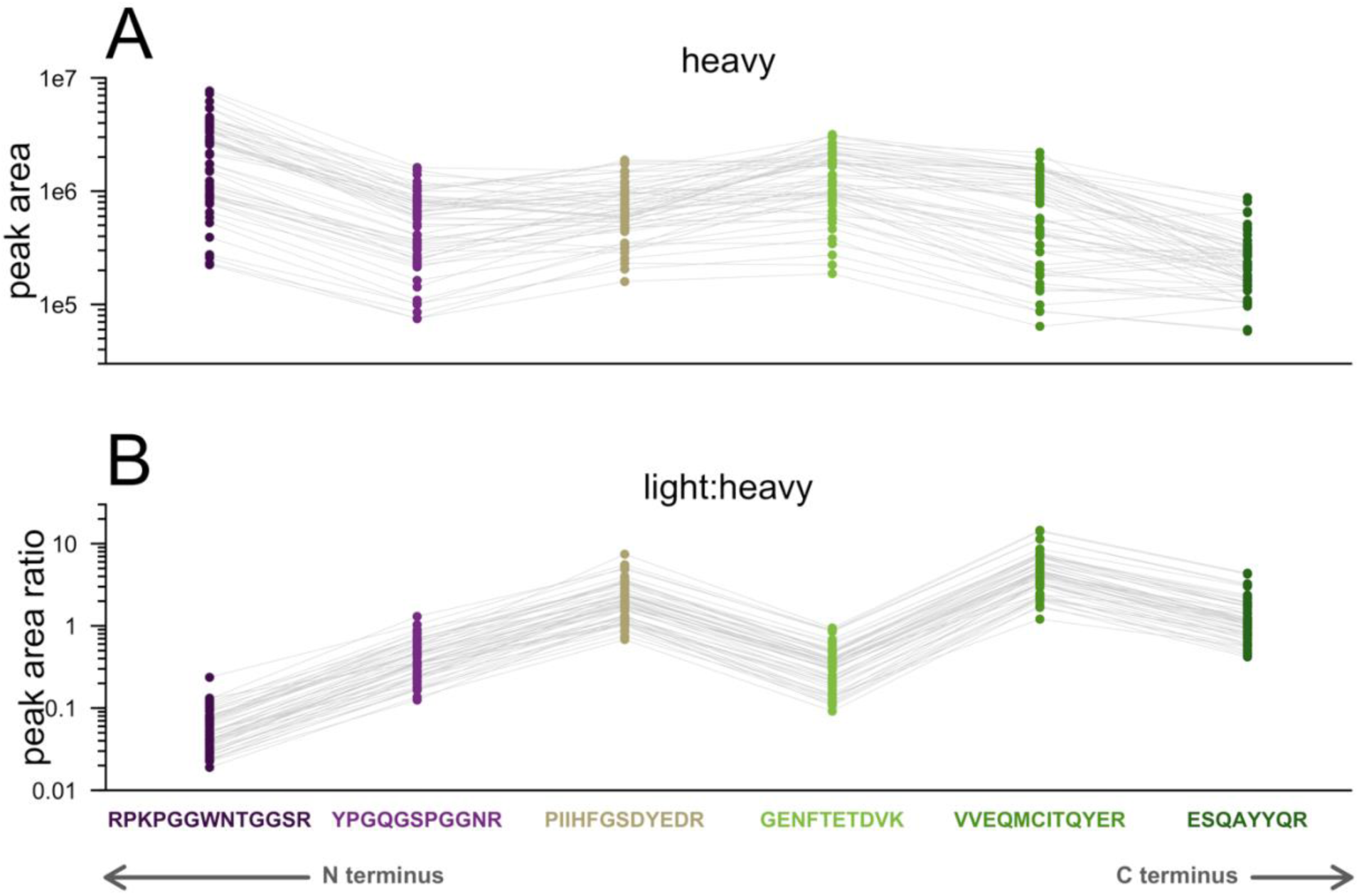
Relative recovery of synthetic heavy peptides in CSF. This plot is the same as Figure 2 but showing **A)** heavy peak area and **B)** light:heavy ratio across the 55 clinical samples.

**Figure S7.**
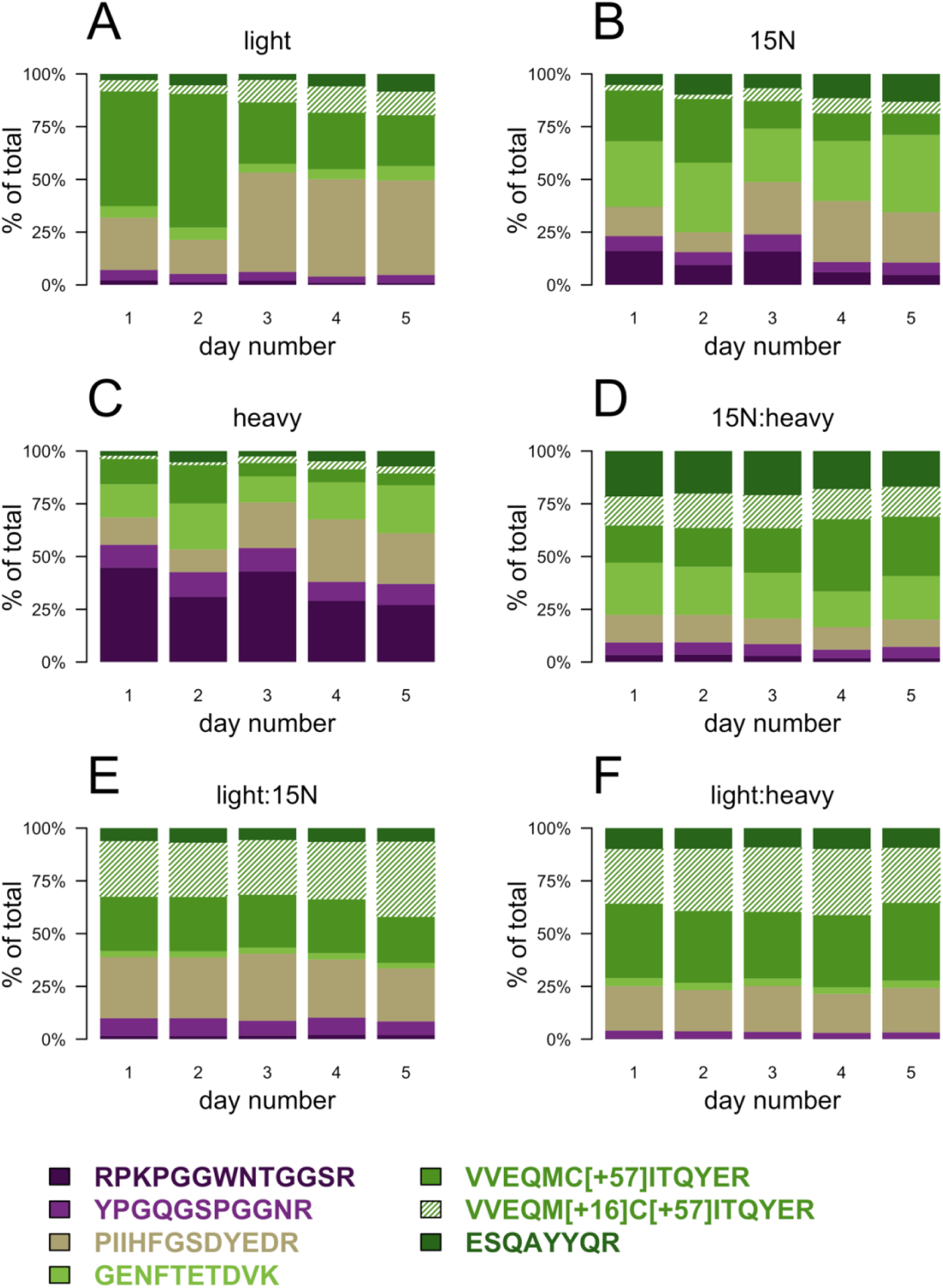
Peptide abundance and ratios by day. Each stacked barplot shows the percent of total PrP peptide abundance contributed by each peptide in clinical samples across five different days.

**Figure S8.**
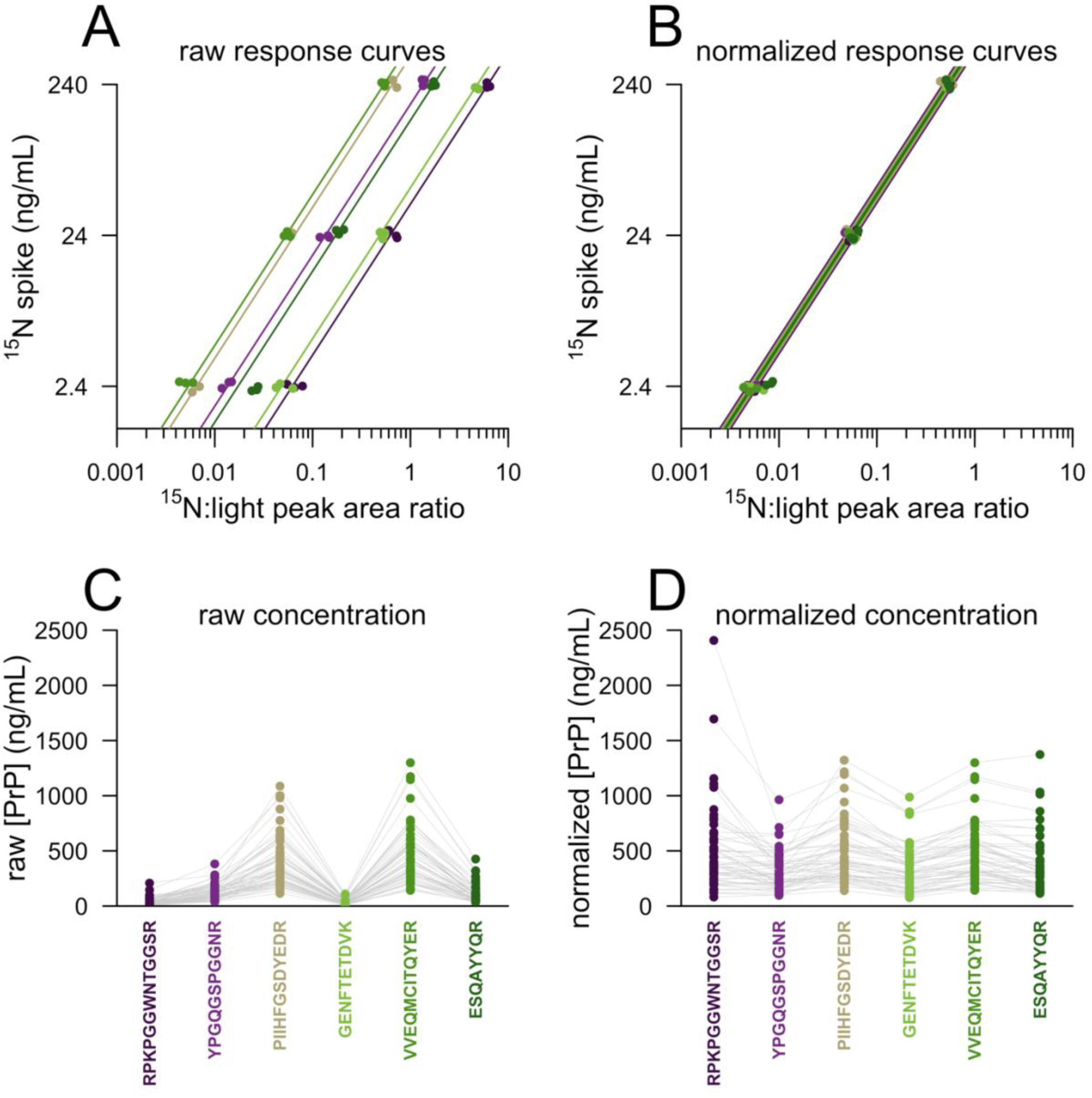
Normalization based on dose-response data. Normalization was performed as described in Methods using data from the same ^15^N dose-response experiment depicted in Figure S5C. **A)** Spiked ^15^N PrP concentration versus observed ^15^N:L ratio for each peptide. Each point is one replicate, and points are jittered along the y axis so that each point is visible. We fit linear models correlating spike ∼ ^15^N:L ratio with the y-intercept fixed at zero, and each peptide yielded a different slope. Note that this figure is plotted in log-log space, so the different slopes appear as different intercepts. We assigned each peptide a response factor equal to the maximum observed slope (that for VVEQMITQYER, top left) divided by its own slope. **B)** Same data from panel A but with response factors applied. **C)** Raw PrP concentrations in clinical samples (simply ^15^N:L ratio times the known ^15^N concentration of 24 ng/mL). **D)** Normalized PrP concentrations in clinical samples (^15^N:L ratio times 24 ng/mL times peptide response factor). In C and D, gray lines connect the dots representing distinct peptides from the same sample.

**Figure S9.**
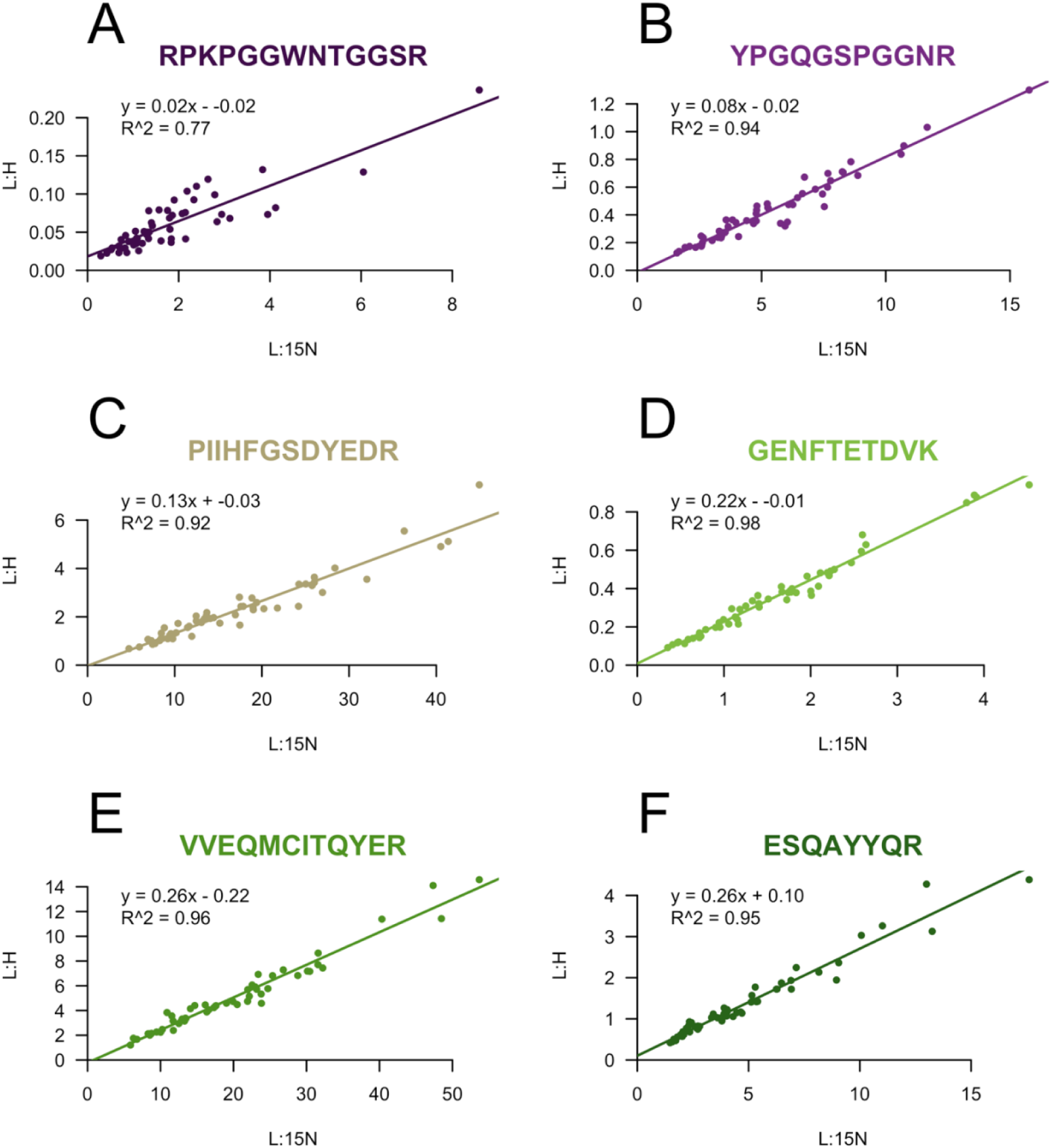
Correlation between L:^15^N and L:H ratios across clinical samples, for each peptide. The correlations exhibit different slopes, consistent with the different observed area of heavy vs. ^15^N across peptides (Figure 2 and Figure S6) and likely arising from differences in recovery from trypsin digestion or other analytical factors prior to heavy peptide addition. Nonetheless, all correlations show good linearity (R^2^ ≥ 0.77), indicating that the two normalization approaches — using synthetic heavy peptides or uniformly ^15^N-labeled recombinant — give similar results.

**Figure S10.**
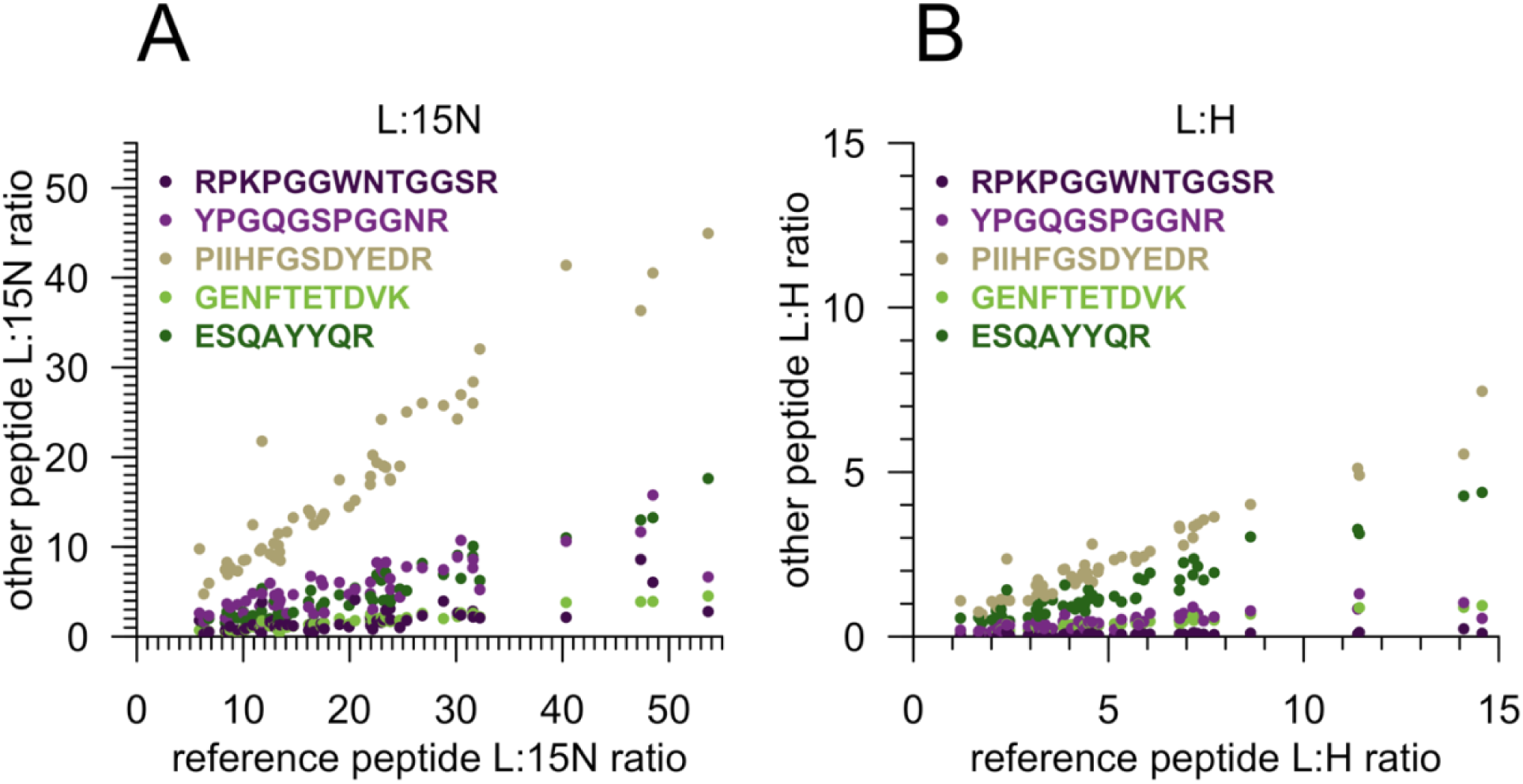
Scatterplot correlations between peptides across clinical samples. **A)** L:^15^N ratio and **B)** L:H ratio. The most abundant peptide, VVEQMCITQYER, is used as reference (x axis) versus all other peptides (y axis). The slopes differ, consistent with different response of different peptides (Table S5 and Figure 2 and S5), but linear correlations are observed for each, across the full dynamic range of samples analyzed. This provides supporting evidence that our assay is technically able to measure the biological variability among samples, and that all peptides move together according to changes in disease state.

